# TGFβ signaling in cancer-associated fibroblasts drives a hepatic gp130-dependent pro-metastatic inflammatory program in CMS4 colorectal cancer subtype

**DOI:** 10.1101/2024.06.25.600311

**Authors:** TJ Harryvan, S Abudukelimu, I Stouten, EJ van der Wel, SGT Janson, KJ Lenos, TRM Lannagan, M White, OJ Sansom, EME Verdegaal, LJAC Hawinkels

**Author notes:** Contributed equally (TJ Harryvan, S Abudukelimu, EME Verdegaal, LJAC Hawinkels). Corresponding author: Dr. Lukas J.A.C. Hawinkels, Department of Gastroenterology and Hepatology, Leiden University Medical Center Albinusdreef 2, 2333 ZA Leiden, The Netherlands.

## Abstract

Current molecular classification of colorectal cancer (CRC), the Consensus Molecular Subtypes (CMS), has highlighted the biological heterogeneity of CRC and enables patient stratification based on the molecular subtype of their tumor. The CMS4 subtype shows the worst prognosis and is linked to the highest occurrence of hepatic metastasis but the underlying molecular mechanisms remain unclear. In this study, we show that the molecular features that largely define CMS4 classification, i.e. abundance of cancer-associated fibroblasts (CAFs) in the tumor microenvironment (TME) and active TGFβ signaling, converge to promote liver metastasis. Studying TGFβ signaling in CRC patient-derived CAFs from the primary tumor revealed that all three TGFβ isoforms induce expression of different IL-6 cytokine family members, particularly IL-6 and IL-11. This primary tumor-derived IL-6 and IL-11 in turn induce upregulation of myeloid chemoattractants, including SAA1, in hepatocytes. Chemical inhibition and genetic ablation experiments revealed that gp130, the IL-6 family of cytokine co-receptor, through JAK/STAT signaling is crucial for the induction of neutrophil chemoattractants by hepatocytes and mediates the migration of potential pro-metastatic neutrophils towards the liver. This IL-6 family-JAK/STAT stromal signaling axis is active in both a murine model of CMS4 as well as in human CRC patients *in vivo*. Combined, our data reveal that TGFβ signaling in CAFs actively contributes to the formation of a neutrophil-dependent, pre-metastatic hepatic niche and that this mechanism might play a role in the metastatic phenotype of CMS4 subtype CRC.

## Introduction

The incidence of colorectal cancer (CRC) is rising, making it one of the most frequently diagnosed malignancies worldwide^1^. The leading cause of death in CRC is the occurrence of distant metastasis, most notably in the liver^2^. Venous drainage of the majority of the colorectum via the portal vein is thought to, at least partly, underlie this hepatic tropism. However, current molecular classifications of primary CRC show that particular CRC subgroups have an increased risk of liver metastasis, indicating that other, tumor-intrinsic factors also modulate metastatic spread to the liver.

The Consensus Molecular Subtypes (CMS) classification was devised to shed light on these tumor-intrinsic factors by means of whole-tumor bulk RNA sequencing^3^. This molecular classification system thus entails both the tumor cells, as well as the adjacent tumor microenvironment (TME), which has been shown to play a crucial role in CRC progression and patient prognosis^4, 5^. Cancer-associated fibroblasts (CAFs) are the most abundant stromal cell subtype in the TME and their abundance is associated with poor prognosis in CRC patients^6–9^. This is also reflected in the CMS classification, in which the CMS4 subtype, comprising 23% of all primary CRCs and characterized by a stromal gene signature^3, 10^.

Current evidence for the aggressive, metastatic phenotype observed in CMS4 tumors points to the Transforming Growth Factor beta (TGFβ) signaling pathway, which is highly activated in CMS4-CRC^3^. TGFβ levels increase during the progression of CRC^11^, and initially restrain tumor growth by inhibiting the proliferation of epithelial tumor cells^12^. In advanced stages of CRC, mutations in genes encoding downstream effectors of canonical TGFβ signaling, such as SMAD4 and TGFβ receptor-II (TβRII)), alleviate the suppressive effects of TGFβ on tumor cell proliferation. However, CAFs and other stromal cell subtypes in the TME retain functional TGFβ signaling^13^, potentially leading to a pro-tumorigenic environment that promotes metastasis initiation^14^ and suppresses or excludes tumor-specific T cells^15–17^.

Whether these two features of the CMS4 subtype, i.e. the high abundance of CAFs and activation of TGFβ signaling, also directly influence hepatic metastasis formation is currently unknown. Insights from other malignancies, particularly pancreatic ductal adenocarcinoma (PDAC)^18, 19^, have highlighted the active modulation of the liver to form a pre-metastatic niche that promotes subsequent tumor seeding from the primary site. In part, this is mediated through influx of pro-metastatic neutrophils, a myeloid cell type that is known to be actively involved in shaping the metastatic niche^20–23^. However, it is currently unknown whether similar mechanisms also contribute to the increased occurrence of hepatic metastasis in CMS4 CRC patients.

In this study, we investigate whether and how TGFβ signaling in CAFs leads to the formation of a hepatic pre-metastatic niche for CRC metastasis. We found that all of the three existing TGFβ isoforms induce the expression of different interleukin-6 (IL-6) cytokine family members, particularly IL-6, and IL-11, in primary patient-derived CRC-CAFs. These cytokines, in turn, trigger IL-6 family cytokine co-receptor gp130-dependent pro-inflammatory response via JAK/STAT signaling in hepatocytes, upregulating neutrophil chemoattractants, most notably Serum Amyloid A1 (SAA1), and promoting neutrophil migration towards the liver. Furthermore, we observed the activity of this gp130 dependent-JAK-STAT signaling axis in human CRC patients and a murine model of CMS4-CRC. Overall, our findings support a pro-metastatic cascade initiated by TGFβ signaling in CRC CAFs, resulting in neutrophil migration to the liver via gp130-mediated signaling in hepatocytes, thereby promoting the increased hepatic metastasis rate observed in CMS4 CRC.

## Materials & methods

### Primary fibroblast isolation, cell culture and conditioned medium (CM)

Normal colon, CRC and liver tissue as well as patient-derived serum were obtained according to the Code of Conduct for Responsible Use of human tissues with approval of the local biobank committee (study ID: B20.006). Primary fibroblasts were derived from human primary and liver-metastasized CRC according to the Code of Conduct for Responsible Use of human tissues or after written informed consent was obtained. Adjacent normal colon or liver tissue was used for the isolation of matched, normal fibroblasts from primary and liver-metastasized CRC resection specimens, respectively. Fibroblasts were isolated and their identity was verified via qPCR as described previously^24^.

Primary fibroblasts and the normal colon myofibroblast cell line CCD-18Co (ATCC) were cultured in Dulbecco’s modified Eagle’s medium (DMEM)/F12 (Thermo Fisher Scientific, Waltham, MA, USA) supplemented with 8% FCS, 100 IU/mL penicillin and 100 μg/mL streptomycin (all Thermo Fisher Scientific). The Huh-7 hepatocyte cell line was obtained from ATCC and cultured in DMEM (4.5g/L D-Glucose) supplemented with 8% FCS, 100 IU/mL penicillin and 100 μg/mL streptomycin (all Thermo Fisher Scientific). The Huh-7 cell line was authenticated and tested negative for mycoplasma contamination. Huh-7 spheroid formation was induced by seeding 3,000 cells in DMEM medium supplemented with 0.24% methylcellulose (Sigma-Aldrich, Zwijndrecht, The Netherlands) in a U-bottom 96-well plate (Greiner bio-one, Frickenhausen, Germany). After 24 hours of incubation, spheroids were transferred to an ultra-low attachment plate (Corning, NY, US) for stimulation and after 30 minutes were washed and processed for immunohistochemistry.

Fibroblast-conditioned medium was acquired from subconfluent cells, either unstimulated or stimulated with 5 ng/ml TGFβ1, TGFβ2, or TGFβ3 for 6 hours, after cells were washed, medium was refreshed (DMEM/F12 0% FCS) and harvested after 48-72 hours. For TGFβ inhibition experiments, the activin receptor like kinase (ALK)-5 kinase inhibitor SB431542 (10 µM, Sigma-Aldrich) was added to the CAFs for the duration of the experiments. For obtaining the Huh-7 conditioned medium, cells were incubated with either unstimulated or TGFβ1 stimulated fibroblast-conditioned medium for 48 hours. All conditioned media were centrifuged to remove cellular debris, aliquoted and stored at - 20 °C until use.

### Constructs, lentiviral transduction, generation and validation of transgenic cell lines

For all lentiviral constructs, third-generation packaging vectors and human embryonic kidney (HEK)293T cells were authenticated and used for the generation of lentiviral particles^25^ in a biosafety level 2 laboratory (BSL-2). To generate a gp130 KO in Huh-7 hepatocytes, a sgRNA, 5’-caccgTGTCAGATCCTTCTCCGCAG-3’ (lowercase nucleotides are compatible with the restriction site) targeting exon 1 was cloned into BsmBI-digested plentiCRISPRv2-puromycin (Add gene: 98290^26^). After transduction, puromycin-resistant Huh-7 cells with no detectable gp130 expression were sorted at single cell density and subsequently expanded to acquire clonal cell lines. Knockout verification of clones was performed via western blot and flow cytometry. Cells were selected and cultured in the presence of 2 µg/ml of puromycin (Sigma).

Rescue experiments were performed by reintroducing a gp130-encoding cDNA in Huh-7 gp130 KO cells, that was amplified from a cDNA library using primers 5’- gatcctcgagaccATGTTGACGTTGCAGACTTG-3’ (forward, XhoI restriction site) and 5’- gatcgctagcTCACTGAGGCATGTAGCCGCC-3’ (reverse, NheI restriction site) and the Phusion High-Fidelity PCR Kit according to the manufacturer’s instruction (Thermo Fisher Scientific). The resulting PCR product was gel purified using the NucleoSpin Gel and PCR Clean-up kit (Macherey-Nagel) and 3’ A-overhangs were added by incubation with Taq*-*polymerase (DreamTaq Green PCR Master Mix (Thermo Fisher Scientific)) for 30 minutes at 72°C. This product was subsequently cloned into the pCR™4-TOPO® TA vector using the TOPO® TA Cloning® kit according to the manufacturer’s instructions (Thermo Fisher Scientific). Subsequently, this clone was sequenced and confirmed to be gp130 transcript variant 1 (NM_002184.4). Finally, this cDNA was subcloned into pLV-CMV-neomycin (kind gift of dr. V. Kemp, dept. Cell & Chemical Biology, LUMC), modified by deletion of the second NheI restriction site present between the IRES and neomycin-resistance gene cassette using site-directed mutagenesis^27^, and designated pLV-THS-CMV-neomycin. The final vector was termed pLV-THS-CMV-gp130-neo. Clonal gp130 KO Huh-7 cells were transduced with this vector and selected and subsequently cultured with neomycin (400 µg/ml; Thermo Fisher Scientific).

The pLV-THS-CMV-gp130-neo vector was used as a template to generate a constitutively active gp130 variant^28^ (gp130^CA^) by deletion of amino acids S187-Y190 in exon 6. To this end, DpnI-mediated site-directed mutagenesis^27^, using primers 5’-cccacctcatgcactgttgattattttgtcaacattgaagtct-3’ (forward) and 5’-agacttcaatgttgacaaaataatcaacagtgcatgaggtggg-3’ (reverse), was performed to obtain the mutant cDNA. Clonal gp130 KO Huh-7 cells were transduced with this mutant vector and selected and subsequently cultured with neomycin (400 µg/ml; Thermo Fisher Scientific). Short hairpin (sh)RNA knockdown constructs were acquired from the Mission TRC shRNA library (Sigma-Aldrich), with target sequences that are shown in supplementary Table 1. Cells were selected and cultured with 2 µg/ml of puromycin (Sigma).

### Real-time quantitative polymerase chain reaction (RT-qPCR^29^)

Total RNA was isolated using the NucleoSpin RNA isolation kit (Macherey-Nagel) according to the manufacturer’s instructions. cDNA was synthesized with the RevertAid First strand cDNA synthesis kit (Thermo Fisher Scientific) using 0.5 – 1.0 µg as RNA input. RT-qPCR was performed with SYBR Green Master mix (Bio-Rad laboratories, Nazareth, Belgium) using the CFX96 Touch Real-Time PCR Detection System (Bio-Rad). Target genes were amplified using specific primers (supplementary Table 2). The ΔCt or ΔΔCt method^30^ was applied to calculate the levels of gene expression, relative to the reference gene (*ACTB*, *HPRT* or *B2M*) or a control condition, respectively. For the TGFβ pathway assays, RT-qPCR was done according to the manufacturer’s instructions (Qiagen, Hilden, Germany).

### IL-6 signaling inhibition

Huh-7 cells were starved overnight in 0% FCS DMEM. The inhibitors anti-IL-6 (siltuximab: Janssen Biologics, Netherlands; 2 µg/mL), anti-IL6R (tocilizumab: Roche, Basel, Switzerland; 8 µg/mL), α-gp130 (R&D Systems, US; 8 µg/mL), Janus kinase (JAK) 1/3 inhibitor (tofacitinib: Pfizer, Netherlands; 5 µM), or human IgG isotype control (Bio X Cell; 2 µg/mL) were added to either TGFβ1-stimulated CAF CM (anti-IL6) or to both starved Huh-7 cells as well as TGFβ1-stimulated CAF CM (anti-IL6R, anti-gp130, JAK inhibitor and human IgG isotype control) for 15 minutes. Subsequently, Huh-7 cells were stimulated with these different mixes for 10 minutes after which cells were harvested.

### Western blot

To investigate the effect of CAF-CM/IL-6 family cytokines induced signaling in hepatocytes, Huh-7 cells were stimulated with undiluted CAF CM or recombinant cytokines for 10 minutes. After stimulation, cells were washed and lysed in radioimmunoprecipitation assay buffer (RIPA) buffer supplemented with Complete® protease inhibitor cocktail (Roche, Basel, Switzerland) and Phosphatase Inhibitor Cocktail II (Abcam, Cambridge, UK) after which protein content was determined using a DC protein assay (Bio-Rad) according to the manufacturer’s instructions. Western blot analysis was performed as described before^24^. Briefly, equal amounts of protein were separated with electrophoresis using 10 or 12% SDS-polyacrylamide gels under reducing conditions. Proteins were transferred to polyvinylidene fluoride (PVDF) membranes (Sigma) and non-specific binding was blocked with 5% milk powder or 5% bovine serum albumin (BSA) in tris-buffered saline containing 0.05% Tween-20 (Merck, Darmstadt, Germany). Blots were incubated overnight with mouse anti-gp130 (R&D systems, Minneapolis, MN, USA, clone 29104), rabbit anti-pSTAT3-Y705 (Cell Signaling Technology, Leiden, The Netherlands, clone D3A7), rabbit anti-STAT3 (Cell Signaling Technology, clone 79D7) or mouse anti-β-actin (Santa Cruz Biotechnology, Dallas, TX, USA, clone C4) as a loading control. Detection was performed by horseradish peroxidase (HRP)-conjugated secondary antibodies (all from Agilent, CA, USA) and chemiluminescence (Thermo Fisher Scientific) was used to visualize the target proteins.

### ELISA

Tissue lysates from (partially) matched primary CRC, liver-metastasis and adjacent normal tissue were lysed in RIPA buffer using the Tissuelyser LT (Qiagen). Total protein content was determined using DC protein assay (Bio-rad) according to the manufacturer’s instructions. IL-6 concentrations in tissue lysates and CAF supernatants were measured using a commercially available kit (Human IL-6 DuoSet ELISA, R&D Systems). SAA1 concentrations in patient sera were measured using a commercially available kit (Human SAA1 DuoSet ELISA, R&D Systems) and a decrease of ≥0.3 OD450 value between post and pre-surgery sera was considered significant. TGF-β1, 2 and 3 levels in tissue lysates and CAF supernatants were determined using an ELISA duoset as described before^31^. Concentrations of cytokines in the tissue lysates were corrected for total protein content.

### Flow cytometry and fluorescence-assisted cell sorting (FACS)

For cell surface staining, Huh-7 cells were harvested and washed twice with FACS buffer, consisting of PBS/0,5% BSA (Sigma) + 0.05% sodium azide (Pharmacy LUMC, Leiden, The Netherlands). Subsequently, cells were incubated with PE-conjugated primary anti-gp130 antibodies (BioLegend, Amsterdam, The Netherlands, clone 2E1 B02) for 45 minutes and washed twice with FACS buffer again. All these FACS experiments were performed on a LSRII (BD Bioscience, CA, USA) and sorting was performed with an Aria FACS sorter (BD Biosciences). Data analyses were performed with FlowJo v10.6.1 (BD biosciences).

### Neutrophil migration assays

Human venous blood was acquired from an in-house voluntary donor biobank (study ID: LuVDS22.016). Neutrophil isolation was performed using the Lymphoprep^TM^ kit (STEMCELL Technologies, Vancouver, Canada) according to the manufacturer’s instructions. To measure neutrophil migration, a Boyden chamber assay was used as described before^32^. Briefly, human neutrophils were labeled with calcein AM (Thermo Fisher Scientific) and left to migrate for 2 hours before termination of the experiment. Fluorescence of calcein-labeled neutrophils was measured on the Cytation Microplate Reader (Biotek, Winooski, VT, USA) at 485 nm and 520 nm wavelength for excitation and emission, respectively.

### KPN organoid generation and intracolonic transplantation

The villinCreER *Kras^G12D/+^Trp53^fl/fl^ Rosa26^N1icd/+^* (KPN) genetically engineered mouse model (GEMM) and tumor-derived organoids were generated as previously described^33^. Briefly, tumors from GEMM mice were cut into small fragments, washed five times in PBS and incubated in 5 ml 10x Trypsin (5mg/ml, Gibco) with 1x DNase buffer and 200U recombinant DNase I (Roche) at 37°C for 30 minutes. After digestion, basal medium was added and the suspension shaken and spun down. The pellet was re-suspended in 10 ml basal medium, filtered through a 70 μm strainer, spun down and pellet suspended in Matrigel (BD Biosciences) according to pellet volume and seeded. Organoids/spheroids were cultured in complete medium at 37°C, 5% CO_2_, 21% O_2_. For culture medium, advanced DMEM/F12 was supplemented with penicillin/streptomycin (100U/ml/100 μg/ml), 2 mM L-Glutamine, 10 mM HEPES, N2-supplement, B27-supplement, 50 ng/ml recombinant mouse EGF and 100 ng/ml recombinant murine Noggin (all from Thermo Fisher Scientific). Organoids were passaged 3 times a week depending on culture confluence.

For intracolonic transplantation, a single male C57BL/6 strain murine KPN organoid line (ID: RBVKPN RKAC3.2f) was used, routinely confirmed to be mycoplasma free before injection. Organoids were harvested and dissociated mechanically, then washed twice with PBS before injection. Approximately 500 organoid fragments in 70 μl PBS were injected per mouse in a single injection. Colonic sub-mucosal injections of organoids were performed as previously described34, using a Karl Storz TELE PACK VET X LED endoscopic video unit. Recipient mice were all male C57BL/6 6-12 weeks old (Charles River, UK) and were monitored until the clinical endpoint, defined by weight loss with or without other body condition changes. Mouse experiments were performed in accordance with national regulatory approval and institutional guidelines.

### Immunohistochemistry

Liver spheroids or tissues were fixed in 4% or 10% neutral buffered formalin, respectively, and processed by standard histology processing techniques as described previously^35^. Ly6G (BioXcell BE0075, 1/60,000 dilution), a marker for murine neutrophils, staining of livers from the intracolonically transplanted mice and WT controls was performed and these were subsequently digitalized using a SCN400F slide scanner (Leica Microsystems, Milton Keynes, UK) at 20x. Scanned images were analyzed using HALO Image analysis software (V2.0.1145, Indica Labs). Whole liver sections were analyzed and results displayed as number of Ly6G positive cells per µm^2^ of liver. Staining of liver spheroids was performed using the rabbit anti-pSTAT3-Y705 (Cell Signaling Technology, clone D3A7) antibody and these were counterstained with hematoxylin.

### RNA sequencing

RNA was isolated using the Qiagen RNeasy Mini kit (Qiagen, 74104) according to the manufacturer’s protocol. Cell pellets or tissue were lysed using the Precellys lysing kit (Bertin Instruments, Montigny-le-Bretonneux, France) in a Precellys Evolution machine (Bertin Instruments). Organoid pellets were snap-frozen and RNA was isolated as described above. RNA of whole tumor samples was isolated at endpoint and conserved in RNAlater (Sigma, R0901) at −80°C until further use. Quality control of isolated RNA and subsequent RNA-sequencing was performed as described previously^33^. Expression levels and differential gene expression were analyzed using DESeq2 package (version 1.24.0) through the Mousr platform^36^.

### Statistical analysis

Data are presented as means ± standard deviation from representative experiments of independent replicates. Unpaired Student t tests were used to compare two independent groups. Paired t-tests were used to compare fibroblasts with or without TGFβ1 stimulation. Differences between more than 2 groups were measured using 1-way analysis of variance (ANOVA) and corrected for multiple testing. All analyses were performed using GraphPad Prism 9.3.1 software (San Diego, CA, USA). P values of 0.05 or less were considered statistically significant.

## Results

### TGFβ1 signaling in primary CRC and LM-CRC CAFs induces expression of IL-6

To identify the genes differentially expressed in primary and liver-metastasis (LM) derived CRC CAFs in a TGFβ1-rich environment, we performed targeted RT-qPCR arrays (**Figure 1A**). Two human primary CRC-CAFs and one LM-CRC CAF were stimulated with TGFβ1 and differentially regulated genes, as compared to unstimulated fibroblasts, were plotted (**Figure 1B**). Upon TGFβ1 stimulation, the majority of TGFβ1 target genes were found to be upregulated, among which 7 genes (*BMP6, CDKN2B, GADD45B, IL6, JUNB, RUNX1, SERPINE1*) consistently exhibited upregulation across all three patient-derived CAFs tested (**Figure 1C, D**). Of note, consistent upregulation of *SERPINE1*, a well-established TGFβ1 target gene^31, 37^, shows the validity of this screen.

**Figure 1.**
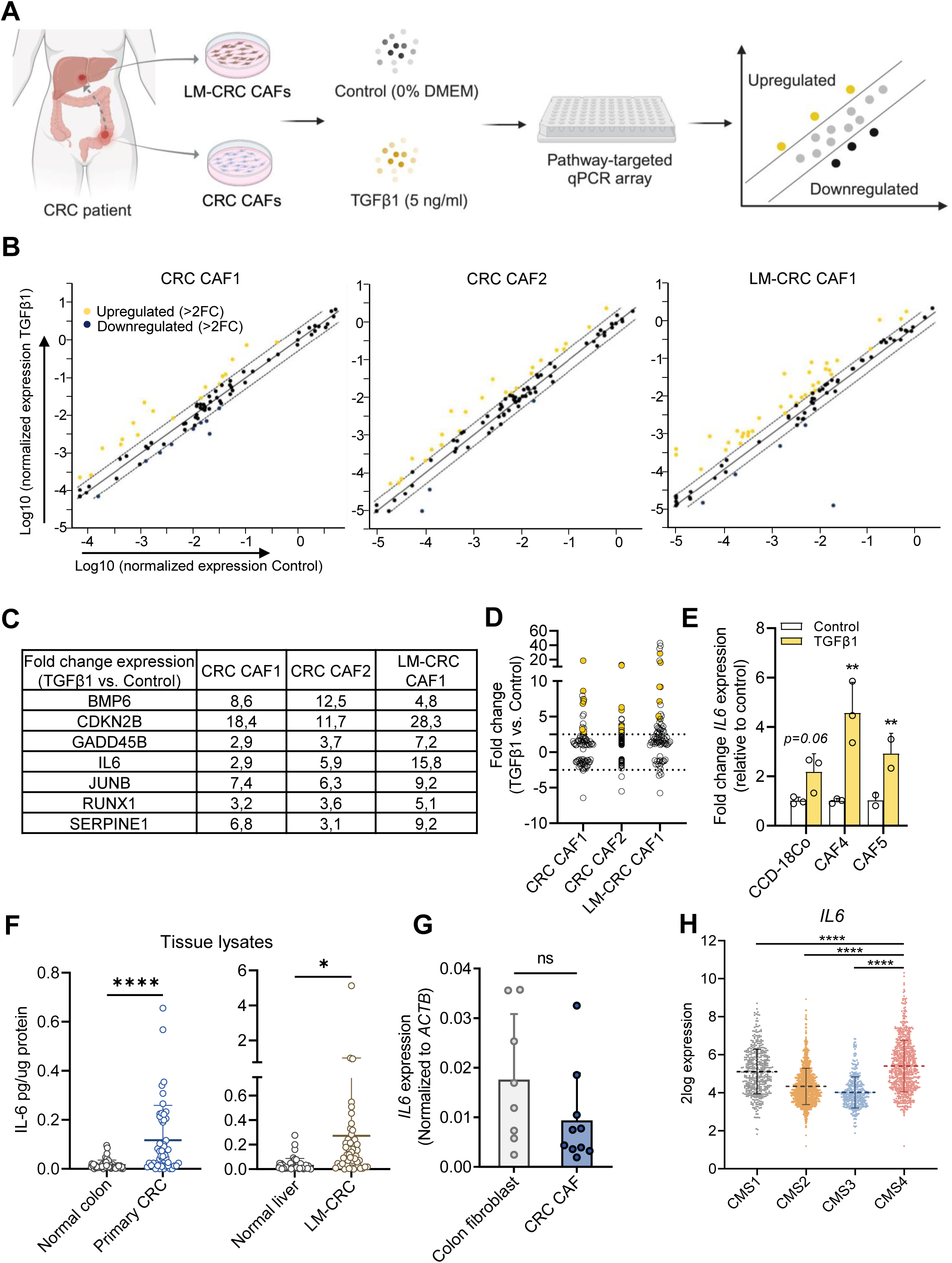
TGFβ1 signaling in primary CRC and LM-CRC CAFs induces expression of IL-6. **A.** Schematic overview of workflow for CAF isolation and TGFβ1 pathway-targeted qPCR array. **B.** RNA expression of TGFβ1 target genes in two primary CRC CAFs (CAF1 and CAF2) and one LM-CRC CAF. qPCR values have been log transformed. **C.** Numerical values depicting the seven TGFβ1 target genes that showed consistent upregulation. **D.** Fold change RNA expression of these seven TGFβ1 target genes (yellow symbols). **E.** *IL6* RNA expression in primary CRC CAFs (CAF4 and CAF5) and the CCD-18Co colon fibroblast line after stimulation with TGFβ1 or medium control. N=3 CCD-18Co, CAF4; N=2 CAF5 independent biological experiments. ** p≤0.01 determined by paired T test. **F.** Protein expression of IL-6 in tissue lysates of normal colon and primary CRC (left panel, N=50) or normal liver and LM-CRC (right panel, N=50) as determined by ELISA. **** p≤0.0001, * p≤0.05 determined by unpaired T test. **G.** *IL6* RNA expression in unstimulated normal colon fibroblasts (N=8) or primary CRC CAFs (N=10). **H.** RNA expression of *IL6* in the original CMS classified cohort^3^ (CMS1: N=457; CMS2: N=1183; CMS3: N=409; CMS4: N=773). **** p≤0.0001 determined by one-way ANOVA with correction for multiple testing (Dunnett’s test).

Two genes encoding for secreted factors, *Bone Morphogenetic Protein (BMP)6* and *IL-6*, were further studied due to their potential paracrine effect on hepatic pre-metastatic niche formation. BMP-6 protein expression was undetectable (all samples <5pg/ml) in TGFβ1-stimulated CAFs, despite reproducible TGFβ1-dependent upregulation at the mRNA level in 2 additional primary CRC-CAFs as well as the normal colon fibroblast cell line CCD-18Co (**supplementary figure 1A**). Moreover, no difference in BMP-6 protein expression was detected between human normal colon and CRC tissues (**supplementary figure 1B**). Therefore this gene was not further investigated. On the contrary, IL-6 was consistently upregulated in two additional independent primary CRC-CAFs upon TGFβ1 stimulation as well as in the CCD-18Co colon fibroblasts (**Figure 1E**). IL-6 protein also showed increased expression was also shown to be increased in tumor tissue in primary and LM-CRC compared to their normal counterpart (**Figure 1F**). These suggest that TGFβ1 induces *IL6* expression in fibroblasts from both normal and malignant colon tissue. Of note, this is only observed in response to TGFβ1 and irrespective of the origin (normal tissue or tumor) of the fibroblasts since we did not observe differential basal expression of *IL6* between normal colon fibroblasts and primary CRC CAFs (**Figure 1G**). This indicates that the TME in which these CAFs reside is important for induction of IL-6 expression. Subsequently, the transcriptional upregulation of *IL6* upon TGFβ1 stimulation was confirmed by cross-referencing our results with an RNAseq dataset (GSE39394^14^) of the normal colon fibroblast cell line CCD-18Co (**supplementary figure 2**).

Next, given the fact that CMS4 subtype of CRC shows a high stromal TGFβ signature, the expression of *IL6* in the different CMS subtypes of a CMS classified cohort^3^ was investigated. Our analysis showed *IL6* to be expressed highest in the CMS4 subtype, compared to all other CMS subtypes (**Figure 1H**). These data show that *IL6* is a bona fide target gene of TGFβ1 signaling in CAFs, with the highest expression in the CMS4 subtype of CRC, characterized by active TGFβ-signaling and an abundance of CAFs.

### IL-6 signaling in CAFs is partly regulated through autocrine TGFβ1 signaling

To further investigate the regulation of *IL6* expression, we examined the basal expression in a cohort of human primary CRC CAFs (N=18) and LM-CRC (N=6) CAFs. LM-CRC CAFs showed higher *IL6* mRNA expression compared to primary CRC CAFs (**Figure 2A**), which was confirmed in an independent publicly available cohort (GSE46824; **Figure 2B**). ELISA analysis confirmed IL-6 in the conditioned medium of these CAFs. Under basal conditions, primary CRC-CAFs and LM-CRC CAFs secreted an average of 280.9 pg/ml and 1157.0 pg/ml of IL-6, respectively (**Figure 2C**). Upon stimulation with TGFβ1, all CAFs showed increased IL-6 production (**Figure 2D, E**) with a similar fold induction between primary and LM-CRC derived CAFs (**Figure 2F**). Considering the disparity in basal *IL6* expression and the strong regulatory effect of TGFβ1, we investigated the involvement of autocrine TGFβ1 signaling in CRC and LM-CRC derived CAFs. Inhibition of the type-I TGFβ receptor activin receptor like kinase-(ALK)5, a crucial downstream mediator, reduced IL-6 secretion to approximately 50% of basal levels in both primary and LM-CRC CAFs (**Figure 2G**). Surprisingly, TGFβ1 RNA (**Figure 2H**) and protein (**Figure 2I**) levels in LM-CRC CAFs did not seem to correlate with their higher IL-6 expression, indicating that additional factors might be involved in regulating IL6 expression. These findings collectively suggest that autocrine signaling via ALK5 is an important mediator of IL-6 expression in CAFs, although additional factors beyond TGFβ1 are probably involved.

**Figure 2.**
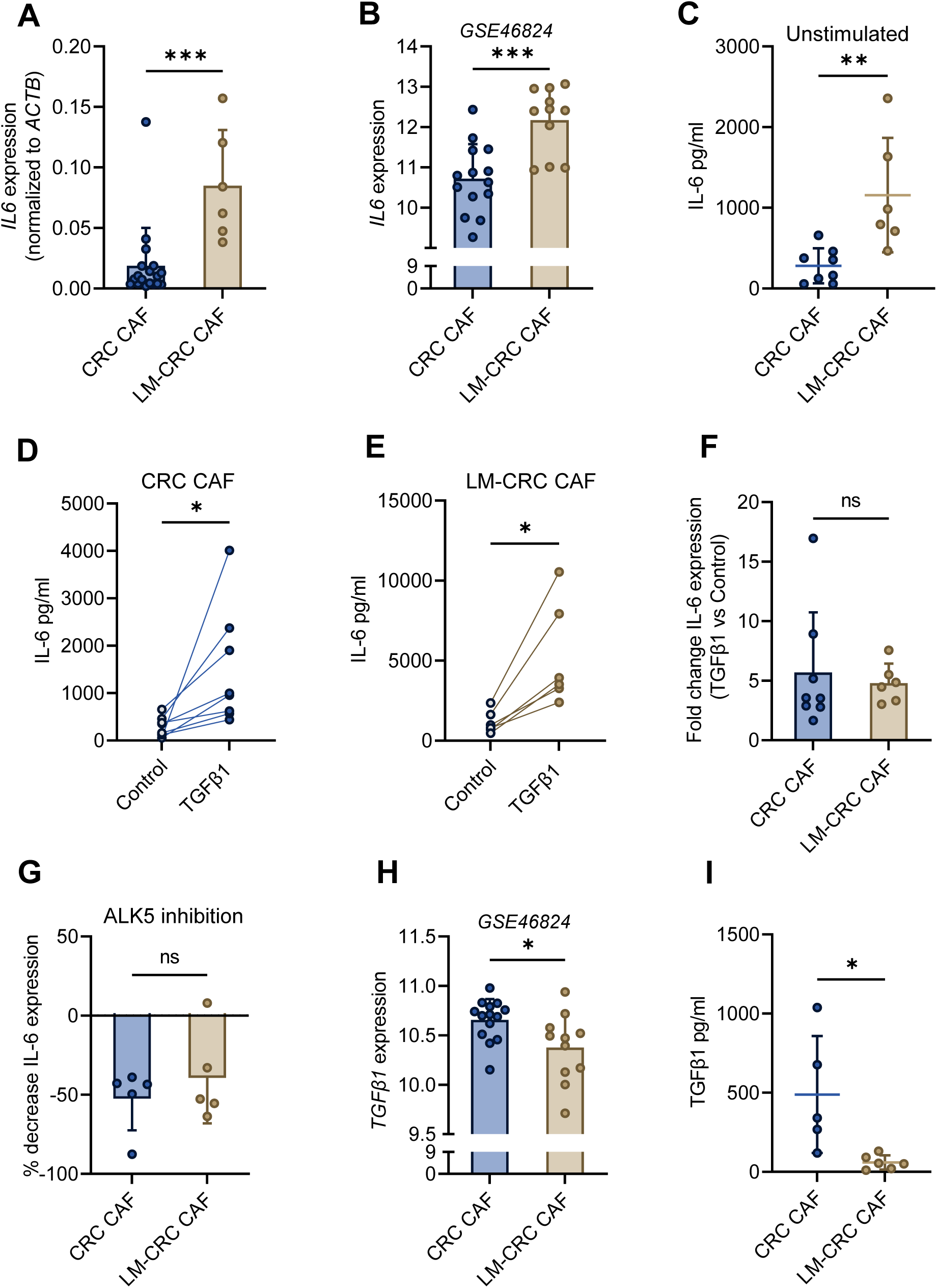
IL-6 signaling in CAFs is partly regulated through autocrine TGFβ1 signaling. **A.** RNA expression of *IL6* in CAFs derived either from the primary tumor (CRC CAF, N=18) or liver metastasis (LM-CRC CAF, N=6). *** p≤0.001 determined by unpaired T test. **B.** RNA-seq data from GSE46824 dataset showing *IL6* RNA expression in an independent cohort of primary CRC CAFs (N=14) and LM-CRC CAFs (N=11). *** p≤0.001 determined by unpaired T test. **C.** IL-6 protein expression in unstimulated primary CRC CAF (N=8) and LM-CRC CAF (N=6) supernatant determined by ELISA. ** p≤0.01 determined by unpaired T test. **D, E.** IL-6 protein expression in TGFβ1-stimulated (5 ng/ml) primary CRC CAFs (N=8) (**D**) and LM-CRC CAFs (N=6) (**E**). * p≤0.05 determined by unpaired T test. **F.** Fold change IL-6 protein expression in supernatant from TGFβ1-stimulated vs. unstimulated primary CRC CAFs (N=8) and LM-CRC CAFs (N=6). ns, p>0.05 determined by unpaired T test. **G.** Percentage decrease in IL-6 protein expression in supernatant in primary CRC (N=5) and LM-CRC (N=5) CAFs after inhibition with SB431532 (Alk5 inhibitor). ns, p>0.05 determined by unpaired T test. **H.** RNA-seq data from GSE46824 dataset (2B) showing *TGFβ1* RNA expression in an independent cohort of primary CRC CAFs (N=14) and LM-CRC CAFs (N=11). * p≤0.05 determined by unpaired T test. **I.** TGFβ1 protein expression in unstimulated primary CRC CAF (N=5) and LM-CRC CAF (N=6) supernatant determined by ELISA. * p≤0.05 determined by unpaired T test.

### All three TGFβ isoforms are elevated in CMS4 CRC and capable of inducing IL-6 expression

The potent impact of ALK5 kinase inhibition on IL-6 expression in LM-CRC CAFs, despite low TGFβ1 expression, led us to investigate the involvement of other TGFβ isoforms signaling through ALK5. Given that TGFβ has three human isoforms (TGFβ1/2/3) with similar or distinct functions depending on the context^38, 39^, we examined their expression in the CMS cohort. Interestingly, all three isoforms were significantly higher expressed in CMS4 CRC compared to CMS1-3 subtypes (**Figure 3A-C**). While TGFβ1 was mainly detected in CRC CAF and not in LM-CRC CAF conditioned medium, TGFβ2 was not detectable in either one and TGFβ3 was expressed at similar levels in both CRC and LM-CRC CAF conditioned medium (**Figure 3D**). This suggests that in primary CAFs, TGFβ1 and TGFβ3, and in LM-CRC CAFs, TGFβ3 may primarily contribute to the autocrine regulation of IL-6 expression. However, considering the extent of IL-6 expression, it is likely that CAFs in the CMS4 TME are exposed to paracrine signals from various cell types expressing different TGFβ isoforms, including tumor cells, macrophages, and endothelial cells^39^.

**Figure 3.**
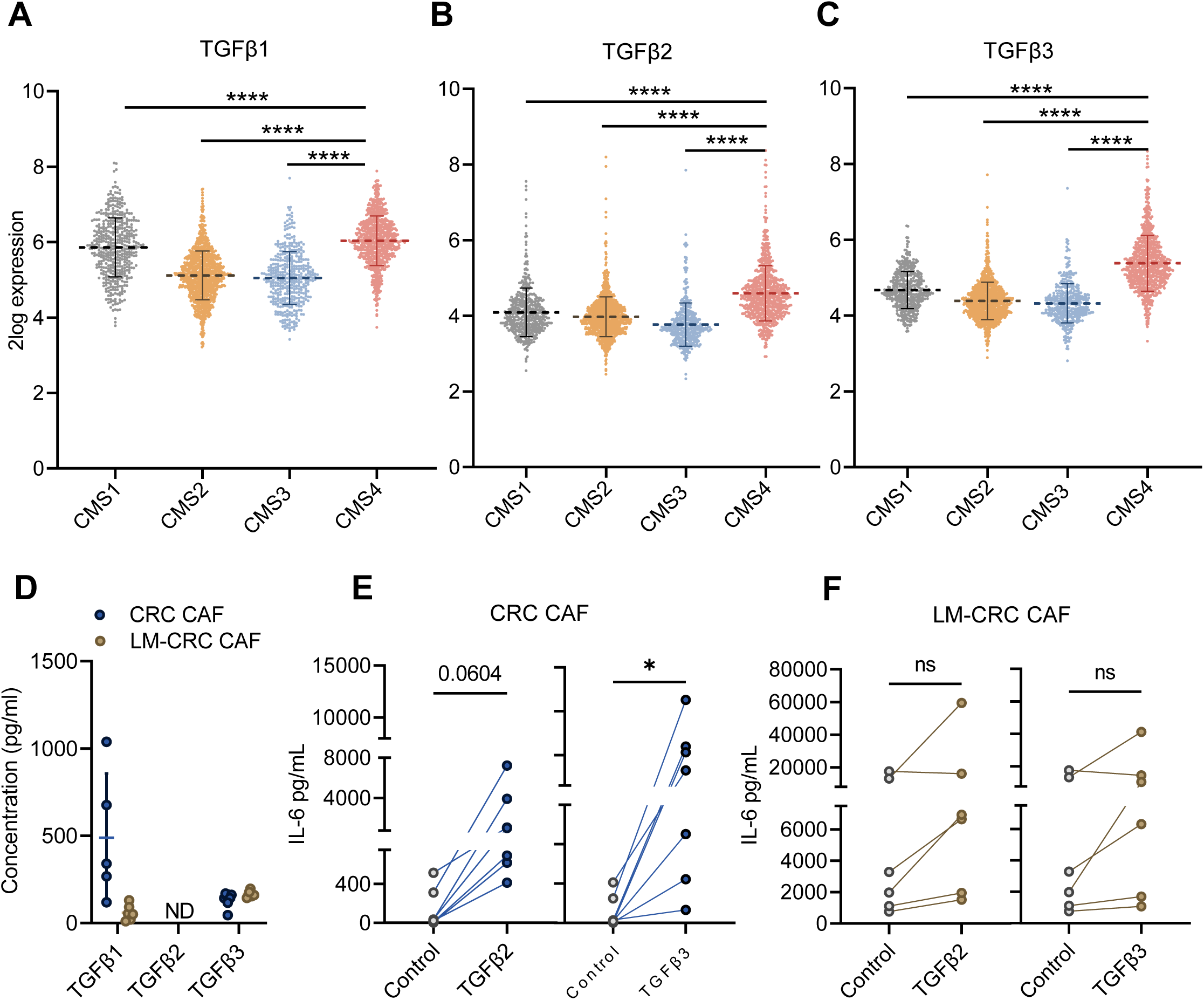
TGFβ isoforms are elevated in CMS4 subtype CRC and capable of inducing IL-6 expression. **A-C.** RNA expression of the three TGFβ isoforms in the original CMS cohort^3^ (CMS1: N=457; CMS2: N=1183; CMS3: N=409; CMS4: N=773). **** p≤0.0001 determined by one-way ANOVA with correction for multiple testing (Dunnett’s test). **D.** Protein expression of TGFβ isoforms in the supernatant of primary CRC and LM-CRC CAFs 48 hours after start of serum starvation as determined by ELISA. **E, F.** Expression of IL-6 in supernatant of primary CRC CAFs (N=7) (**E**) or LM-CRC CAFs (N=6) (**F**) 48 hours after start stimulation with either TGFβ2 or TGFβ3. * p≤0.05 determined by paired T test.

To investigate the effects of exogenous TGFβ2 and TGFβ3 stimulation on IL-6 expression, primary and LM-CRC CAFs were treated with these TGFβ isoforms. Both isoforms induced IL-6 expression in primary CRC as well as LM-CRC CAFs (**Figure 3E, F**). It is worth noting that the expression in LM-CRC CAFs was not significantly upregulated, probably due to already high basal level of IL-6 expression.

In summary, all three TGFβ isoforms are highly expressed in CMS4 CRC and regulate IL-6 expression in both autocrine (TGFβ1, TGFβ3) and paracrine fashion (TGFβ1, TGFβ2, TGFβ3).

### TGFβ-driven IL-6 production by CAFs triggers the expression of neutrophil chemoattractants *SAA1* and *CXCL5* in hepatocytes

Having established the TGFβ-IL-6 signaling axis in CAFs, we next sought to investigate its potential connection to the increased risk of liver metastasis in CMS4 CRC. Clinically, IL-6 has already been linked to CRC progression and the risk of liver metastasis^40^. To understand how TGFβ-IL-6 signaling in primary CRC CAFs influences the formation of a pre-metastatic niche in the liver, we stimulated human Huh-7 hepatocytes with TGFβ1-stimulated CRC CAF conditioned medium (TGFβ1 CAF CM), further referred to as ‘CAF-priming of hepatocytes’ (**Figure 4A**). Stimulation of Huh-7 with TGFβ1 CAF CM from three different CRC patients led to rapid and robust phosphorylation of STAT3 (pSTAT3) compared to unstimulated CRC-CAF CM or control medium (**Figure 4B**). This indicates that TGFβ-CAF CM contains higher levels of cytokines that induce STAT3 phosphorylation. Furthermore, both recombinant IL-6 and TGFβ1 CAF CM significantly increased (P<0.05 and FC>2) the expression of the neutrophil chemoattractants *SAA1*,*CXCL5, and CXCL3* in Huh-7 hepatocytes (**Figure 4C, supplementary figure 3A**), but not that of *CXCL1*, *CXCL2, CXCL6, CXCL7, CXCL8* (**supplementary figure 3A**). This suggests that TGFβ-induced IL-6 expression in the CAF CM is responsible for the upregulation of the neutrophil chemoattractants *SAA1* and *CXCL5* in hepatocytes. To confirm that the IL-6 in TGFβ1 CAF CM is indeed instrumental in regulating *SAA1* and *CXCL5* expression in hepatocytes, we blocked IL-6 signaling at various levels of the signaling cascade (**supplementary figure 3B**). Direct blocking of IL-6 in TGFβ1 CAF CM (anti-IL-6; siltuximab) or its receptor (anti-IL6R; tocilizumab) partially reduced the upregulation of *SAA1* and *CXCL5* compared to an isotype control (**Figure 4D, E**), while inhibiting downstream signaling by blocking the IL-6 co-receptor gp130 (α-gp130) or Janus kinase (JAK) 1/3 (tofacitinib), a clinically approved kinase inhibitor, completely abolished the induction of *SAA1* and *CXCL5* in CAF-primed Huh-7 (**Figure 4D,E**).

**Figure 4.**
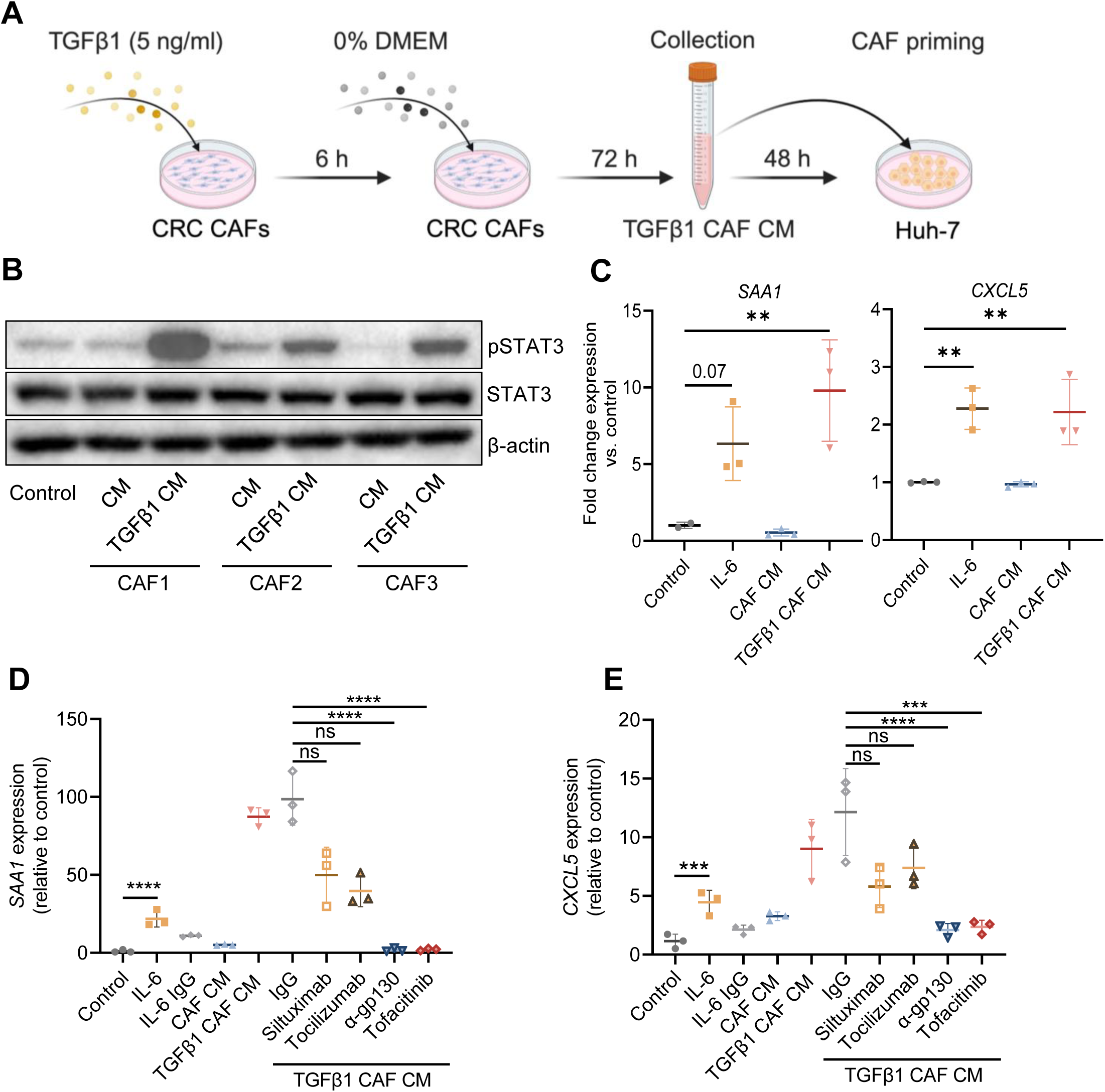
TGFβ-mediated IL-6 production by CAFs induces the expression of neutrophil chemoattractants *SAA1* and *CXCL5* in hepatocytes. **A.** Schematic overview of ‘CAF priming of Huh-7 hepatocytes’, referring to Huh-7 cells are exposed to TGFβ1-stimulated CRC CAF CM **B.** Western blot for pSTAT3 and total STAT3 in wildtype Huh-7 cells that were stimulated with CAF CM or TGFβ1 CAF CM from 3 different CRC CAF lines. **C.** RNA expression of *SAA1* and *CXCL5* in wildtype Huh-7 cells stimulated with DMEM 0% FCS (Control), IL-6 (50 ng/ml), CAF CM or TGFβ1 CAF CM. N=2 or N=3 for the Control group, due to the very low *SAA1* expression; for the other conditions, N=3.*** p≤0.001, ** p≤0.01, * p≤0.05 determined by one-way ANOVA with correction for multiple testing (Dunnett’s test). **D, E** RNA expression of *SAA1* and *CXCL5* in wildtype Huh-7 cells after CAF priming with blockade of specific components of the IL-6 signaling pathway. Siltuximab (anti-IL-6, 2 µg/mL), tocilizumab (anti-IL6R, 8 µg/mL), α-gp130 (anti-gp130, 8 µg/mL), tofacitinib (anti-JAK1/3, 5 µM), or human IgG isotype control (Bio X Cell; 2 µg/mL) were added to Huh-7 cells along with TGFβ1-stimulated CAF CM for 15 minutes. Subsequently, these different stimulations were applied to Huh-7 cells for 10 minutes. N=1 for all different conditions. **** p≤0.0001, *** p≤0.001 determined by one-way ANOVA with correction for multiple testing (Dunnett’s test).

Overall, these data suggest that TGFβ-IL-6 signaling in primary CRC CAFs contributes to the induction of the neutrophil chemoattractants *SAA1* and *CXCL5* in hepatocytes and blocking IL-6 signaling results in a partial reduction in their upregulation.

### The induction of neutrophil chemoattractants in hepatocytes is fully gp130-dependent

We next explored whether the induction of neutrophil chemoattractants in hepatocytes is dependent on gp130. In line with the complete abrogation of chemoattractant induction by the gp130 blocking antibody, α-gp130 also entirely inhibited STAT3 phosphorylation in CAF-primed Huh-7 hepatocytes (**Figure 5A**). To further determine the critical role of gp130 in hepatocytes, shRNA-mediated knockdown of gp130 was performed and showed similar effects (**supplementary figure 4A-C)**, although the signaling was not completely inhibited due to residual gp130 expression. Therefore, to validate these findings, a clonal Huh-7 gp130 KO was generated using CRISPR/Cas9 (**supplementary figure 4D**), and deletion of gp130 in Huh-7 hepatocytes completely abolished STAT3 phosphorylation induced by TGFβ1 CAF CM (**Figure 5B**). Moreover, the gp130 KO cell line was rescued by reintroducing gp130 cDNA (**supplementary figure 4E**), which fully restored pSTAT3 induction upon stimulation with TGFβ1 CAF CM (**Figure 5B**). Notably, the downstream *SAA1* induction by TGFβ1-primed CAF CM was only observed in Huh-7 cells with intact gp130 signaling (**Figure 5C**). Finally, CAF-priming of hepatocytes was visualized by pSTAT3 staining of Huh-7 spheroids with and without intact gp130 signaling, further confirming the indispensable role of gp130 in CAF-induced STAT3 phosphorylation in hepatocytes (**Figure 5D**).

**Figure 5.**
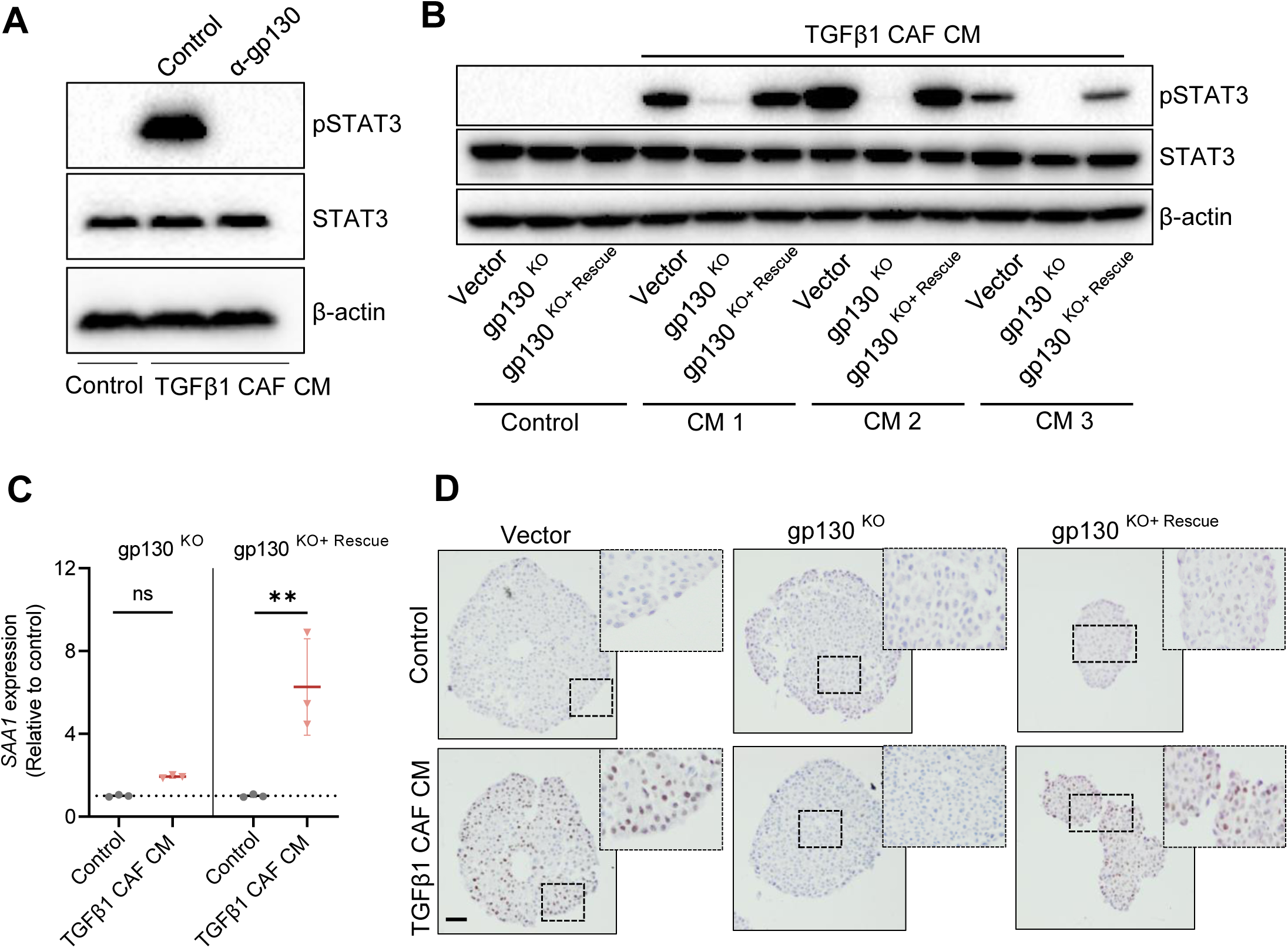
The induction of neutrophil chemoattractants in hepatocytes is fully gp130-dependent. **A.** Western blot for pSTAT3 and total STAT3 in CAF CM-primed Huh-7 cells in which gp-130 signaling is blocked using α-gp130 antibody. α-gp130, anti-gp130. β-actin is used as a loading control. **B.** Western blot for pSTAT3 and total STAT3 in vector control, gp130^KO^, or gp130^KO^ rescued Huh-7 cells (gp130^KO+Rescue^) after stimulation with DMEM 0% (Control) or TGFβ1 CAF CM. **C.** RNA expression of *SAA1* in gp130^KO^ and gp130^KO+Rescue^ Huh-7 cells after CAF priming (N=3 independent biological experiments). ** p≤0.01 determined by unpaired T test. **D.** IHC of pSTAT3 in Huh-7 cells harboring a vector control, gp130^KO^ or gp130^KO+Rescue^ after stimulation with DMEM 0% or after CAF priming. Scale bar, 50µm.

As specific inhibition of IL-6 only partially blocked STAT3 and downstream target genes, these data indicate that, in addition to IL-6, other CAF-derived proteins requiring gp130 signaling that are induced by TGFβ1 are involved in the pro-inflammatory program induced by CAF-priming of hepatocytes. Therefore, we examined the expression of the 7 other members of the IL-6 cytokine family that utilize gp130 as a co-receptor in CRC CAFs stimulated with TGFβ1 (**supplementary figure 5A**). Alongside *IL6*, *IL11* and *LIF* showed strong upregulation in at least 4 out of 5 primary CRC CAFs tested (**supplementary figure 5B**), indicating their potential contribution to pSTAT3 induction and upregulation of neutrophil chemoattractants. Upregulation of *IL11* and *LIF* in CCD-18Co colonic fibroblasts after TGFβ1 stimulation was validated in an external cohort (GSE39394^14^; **supplementary figure 2**). To further substantiate the role of IL-11 in pSTAT3 induction in CAF-primed Huh-7, shRNA-mediated knockdown of the IL-11 receptor (IL11R) was performed (**supplementary figure 5C**). Knockdown of the IL11R on Huh-7 hepatocytes resulted in a strong reduction in STAT3 phosphorylation upon CAF-priming of hepatocytes (**supplementary figure 5D**), further confirming the role of IL-11 in the CAF-mediated priming of hepatocytes.

Taken together, these results suggest that IL-6, IL-11 and LIF are induced by TGFβ1 signaling in CAFs and that CAF-derived IL-6 and IL-11 are capable of inducing pSTAT3 in hepatocytes in a gp130-dependent fashion.

### Hepatic gp130 signaling regulates neutrophil migration

Since neutrophils are described to play an important role in (pre-)metastatic niche formation^20–22^, including the liver^19, 23^, we argued that CAF-driven gp130-signaling in hepatocytes might contribute significantly to the chemotaxis of pro-metastatic neutrophils in CMS4 CRC. To further investigate this hypothesis, we performed neutrophil migration assays using supernatant from CAF-primed Huh-7 hepatocytes (**Figure 6A**). Unstimulated Huh-7 CM only resulted in a small induction of neutrophil migration (**Figure 6B**). However, both TGFβ1-stimulated primary CRC CAF and LM-CRC CAF-primed Huh-7 CM significantly induced neutrophil chemotaxis compared to unstimulated Huh-7 supernatant (**Figure 6B**). Importantly, TGFβ1 CAF CM by itself did not induce neutrophil chemotaxis, indicating that CAF CM did not contain neutrophil chemoattractants. Moreover, stimulation of Huh-7 with TGFβ1 CAF CM resulted in increased neutrophil chemotaxis compared to unstimulated CAF CM (**Figure 6C**). This aligns with the observation that stimulation of Huh-7 with TGFβ1-stimulated CAF CM, but not unstimulated CAF CM, leads to the upregulation of several neutrophil chemoattractants (*SAA1, CXCL5*; **Figure 4C**).

**Figure 6.**
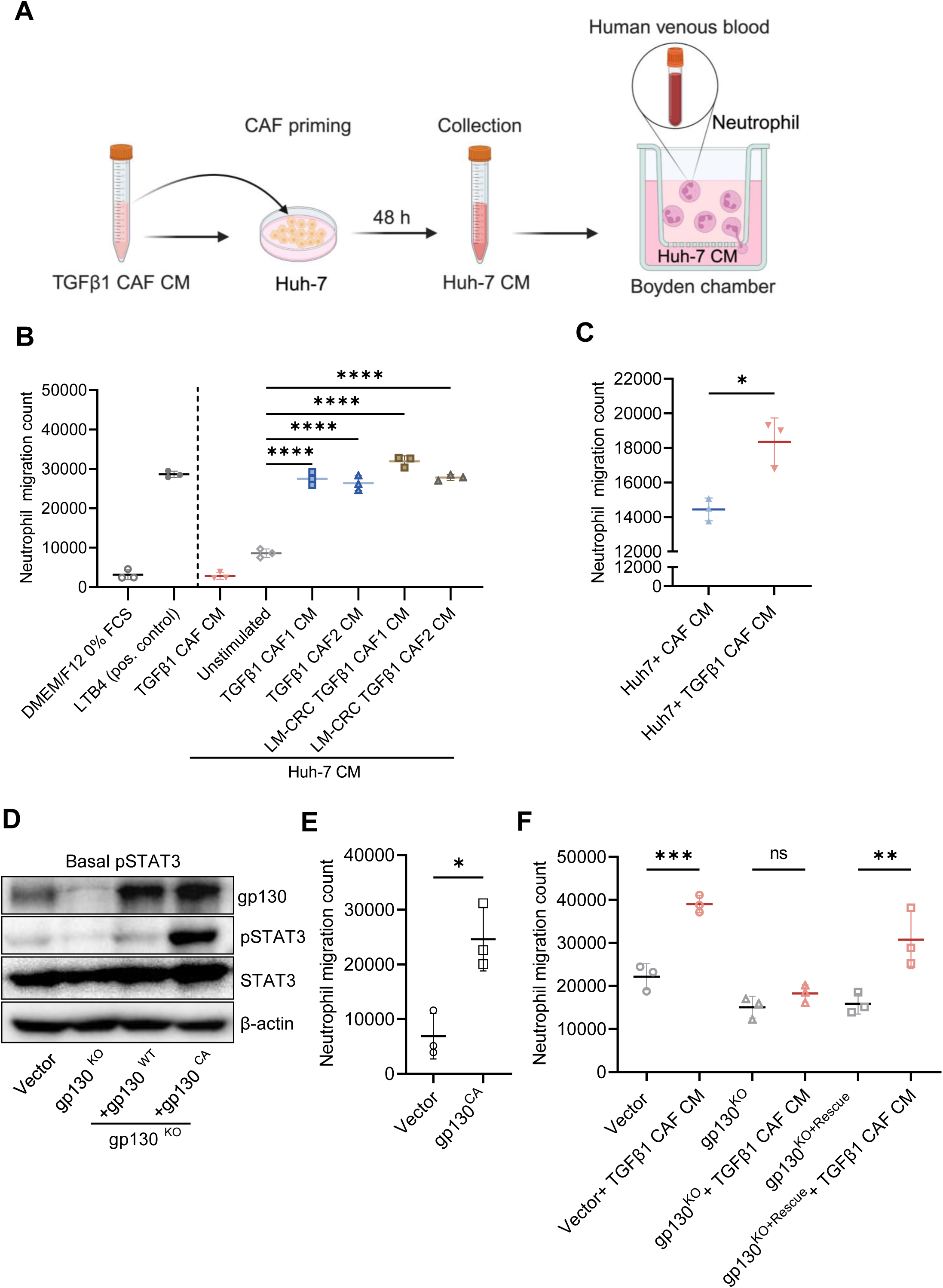
Hepatic gp130 signaling regulates neutrophil migration. **A.** Schematic overview of neutrophil migration assays towards CAF-primed Huh-7 hepatocyte supernatant. **B.** Neutrophil migration count towards supernatant of CAF-primed Huh-7 cells (N=3 independent biological experiments). **** p≤0.0001 determined by one-way ANOVA with correction for multiple testing (Dunnett’s test). **C.** Neutrophil migration (cell counts) towards supernatant of Huh-7 cells stimulated with CAF CM or TGFβ1 CAF CM (‘CAF priming’) (N=3 independent biological experiments). * p≤0.05 determined by unpaired T test. **D.** Western blot for gp130, pSTAT3 and total STAT3 in unstimulated Huh-7 hepatocytes (basal conditions) that contain either vector control, gp130^KO^ or a gp130^KO^ that is rescued with either gp130^WT^ or constitutively active gp130^CA^. β-actin is used as loading control. **E.** Neutrophil migration count towards supernatant of vector control or gp130^CA^ Huh-7 cells (N=3 independent biological experiments). * p≤0.05 determined by unpaired T test. **F.** Neutrophil migration towards unstimulated or CAF-primed Huh-7 cells with vector control, gp130^KO^ or gp130^KO+Rescue^ (N=3 independent biological experiments). *** p≤0.001, ** p≤0.01 determined by one-way ANOVA with correction for multiple testing (Dunnett’s test).

Next, we examined whether the increase in neutrophil chemotaxis was driven by enhanced hepatic gp130-dependent signaling. For this purpose, a constitutively active gp130 variant^28^ (gp130^CA^) was generated and expressed in Huh-7 gp130 KO cells (**supplementary figure 4F**). gp130^CA^ Huh-7 cells, in contrast to gp130 wildtype, have constitutively active gp130 signaling as shown by strong phosphorylation of STAT3 under basal conditions (**Figure 6D**). The supernatant of Huh-7 expressing gp130^CA^, obtained under basal conditions, resulted in enhanced neutrophil migration, confirming that hepatic gp130 signaling is indeed a strong driver of neutrophil chemotaxis (**Figure 6E**).

Finally, to demonstrate that the increased neutrophil chemotaxis observed after CAF-priming of hepatocytes is dependent on gp130 signaling, we used supernatant from Huh-7 cells with or without gp130 KO. CAF-priming of Huh-7 cells with deleted gp130 almost completely abolished the induction of neutrophil chemotaxis, whereas the induction was present in vector control and gp130 KO cells with rescued gp130 expression (**Figure 6F**).

In summary, these data show that hepatic gp130 signaling, activated by CAF-secreted IL-6 cytokine family members, drives neutrophil chemotaxis.

### The IL-6 family of cytokine-JAK-STAT signaling axis is active in CMS4 CRC *in vivo*

We next employed the previously published villinCreER *Kras^G12D/+^ Trp53^fl/fl^ Rosa26^N1icd/+^*genetically engineered mouse model (KPN GEMM^33^) to study the role of this signaling pathway *in vivo.* This GEMM of KRAS^G12D^ driven CRC develops stroma-rich, CMS4 primary tumors with a high penetrance of liver metastasis (80%) that is neutrophil-dependent. Therefore, KPN tumor-derived organoids were used to study the role of this signaling pathway *in vivo* by transplanting them into the colon of immune competent mice. Animals were sacrificed after the development of primary CRC but before the development of liver metastasis to study the formation of a pre-metastatic hepatic niche (**Figure 7A**). mRNA sequencing of primary tumor tissues revealed that, compared to normal colon, there was a strong enrichment of the IL6-JAK-STAT signaling pathway in primary KPN tumors (**Figure 7B**). Additionally, the expression of the individual TGFβ-regulated members of the IL-6 cytokine family, *IL6, IL11* and *LIF*, were significantly higher in KPN tumors compared to normal colon tissue (**Figure 7C**). Notably, the expression of these cytokines also appeared higher in the whole tumor tissue than in the KPN organoids from which the tumors originated (**supplementary figure 7**), indicating that these transcripts are probably stroma derived.

**Figure 7.**
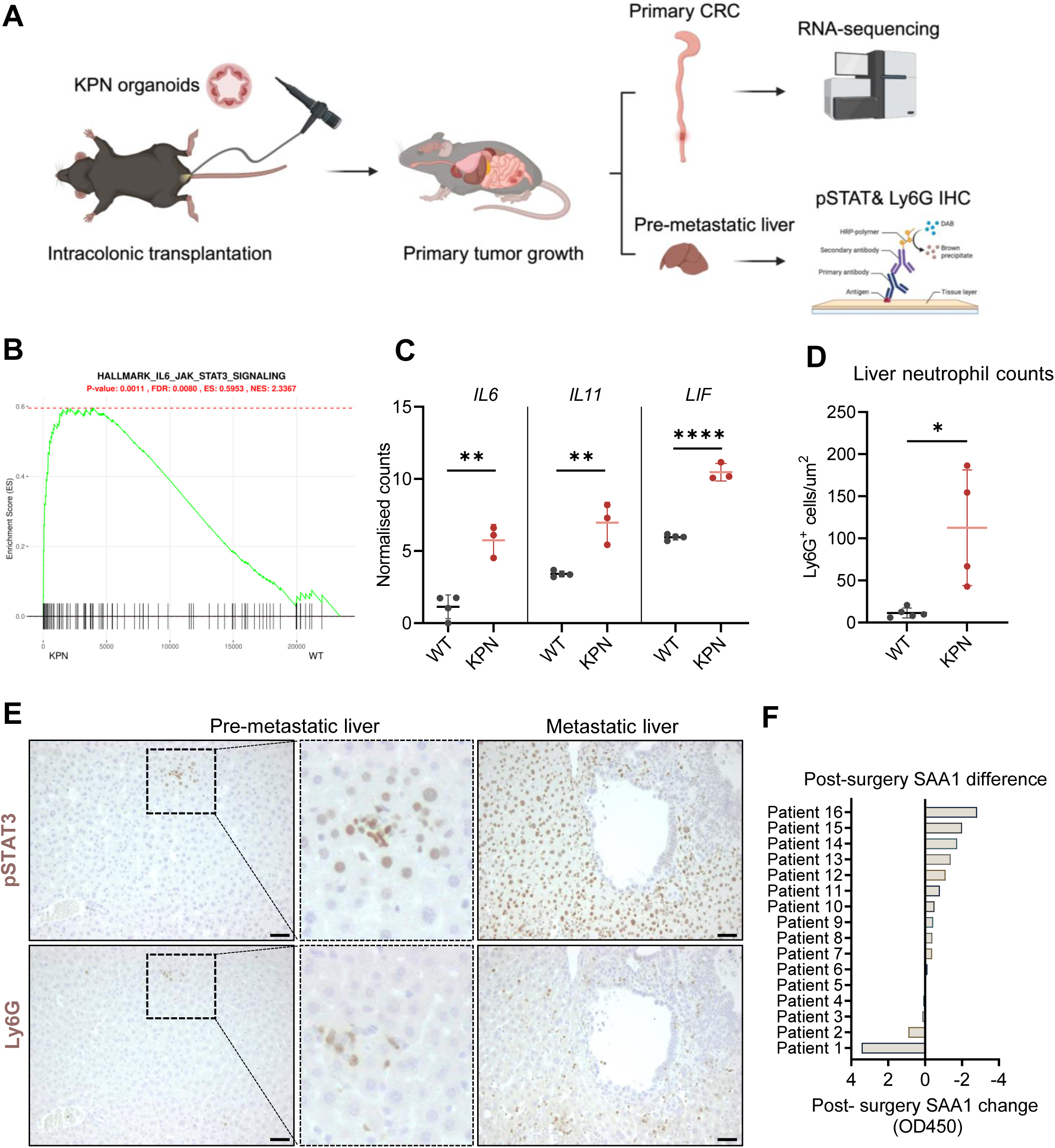
The IL-6 family of cytokine-JAK-STAT signaling axis is active in CMS4 CRC *in vivo*. **A.** Schematic overview of the KPN GEMM organoid experiment. **B.** Gene Set Enrichment analysis showing the Enrichment Score (ES) for the IL6-JAK-STAT signaling hallmark in KPN tumor tissue (N=3) and wildtype normal colon (N=3). **C**. Normalized counts of *IL6, IL11* and *LIF* in KPN tumor tissue (KPN) and normal colon tissue (WT). **** p≤0.0001, ** p≤0.01 determined by unpaired T test. **D.** Quantification of automated scoring of Ly6G staining in pre-metastatic livers of KPN transplanted mice (N=4) and WT livers (N=5). **E.** Representative IHC of pSTAT3 and Ly6G in pre-metastatic and metastatic livers in the KPN GEMM. Scale bar, 50µm. **F.** Waterfall plot of difference in serum SAA1 OD450 value before and after surgery in 16 patients with primary CRC without liver metastasis.

Having established the activation of the IL6-JAK-STAT signaling pathway in this murine model, we investigated the presence of neutrophils in the liver during the pre-metastatic stage. An increased number of Ly6G positive cells in the livers of mice bearing KPN primary tumors compared to non-tumor-bearing WT controls was seen, prior to the development of metastasis (**Figure 7D/E**). Moreover, we observed co-localization of pSTAT3^+^ hepatocytes and Ly6G^+^ neutrophils, although the number of pSTAT3^+^ hepatocytes was low at this stage (**Figure 7E**). In contrast, livers with macroscopic metastases exhibited abundant nuclear pSTAT3 staining (**Figure 7E**). This suggests that the mechanism underlying pre-metastatic niche formation continues to play a significant role even when overt liver metastases are already present. Finally, to assess the clinical relevance of our findings, we measured SAA1 protein levels in the serum of patients with primary CRC but without known liver metastasis at time of diagnosis. SAA1, predominantly produced by hepatocytes, served as a surrogate marker for probing the activity of the IL-6 family of cytokine-JAK-STAT signaling pathway in these patients. By comparing serum SAA1 levels before and after removal of the primary tumor, we could evaluate differences in the activity status of this pathway. Ten out of sixteen (62.5%) CRC patients showed a decrease in serum SAA1 levels three months post-surgery, following resection of the primary tumor **(Figure 7F**), suggesting that SAA1 production by the liver is partly driven by the presence of the primary tumor.

Taken together, we propose a model for hepatic metastasis in CMS4 CRC (**Figure 8**) in which TGFβ drives the production of IL-6 cytokine family members, particularly IL-6 and IL-11, in CAFs. The IL-6 cytokine family induces the activation of hepatocytes, triggering a pro-inflammatory gp130-dependent signaling program that leads to the production of neutrophil chemoattractants. Ultimately, this process results in the recruitment of pro-metastatic neutrophils that aid the formation of a hepatic pre-metastatic niche, allowing subsequent seeding of the primary tumor.

**Figure 8.**
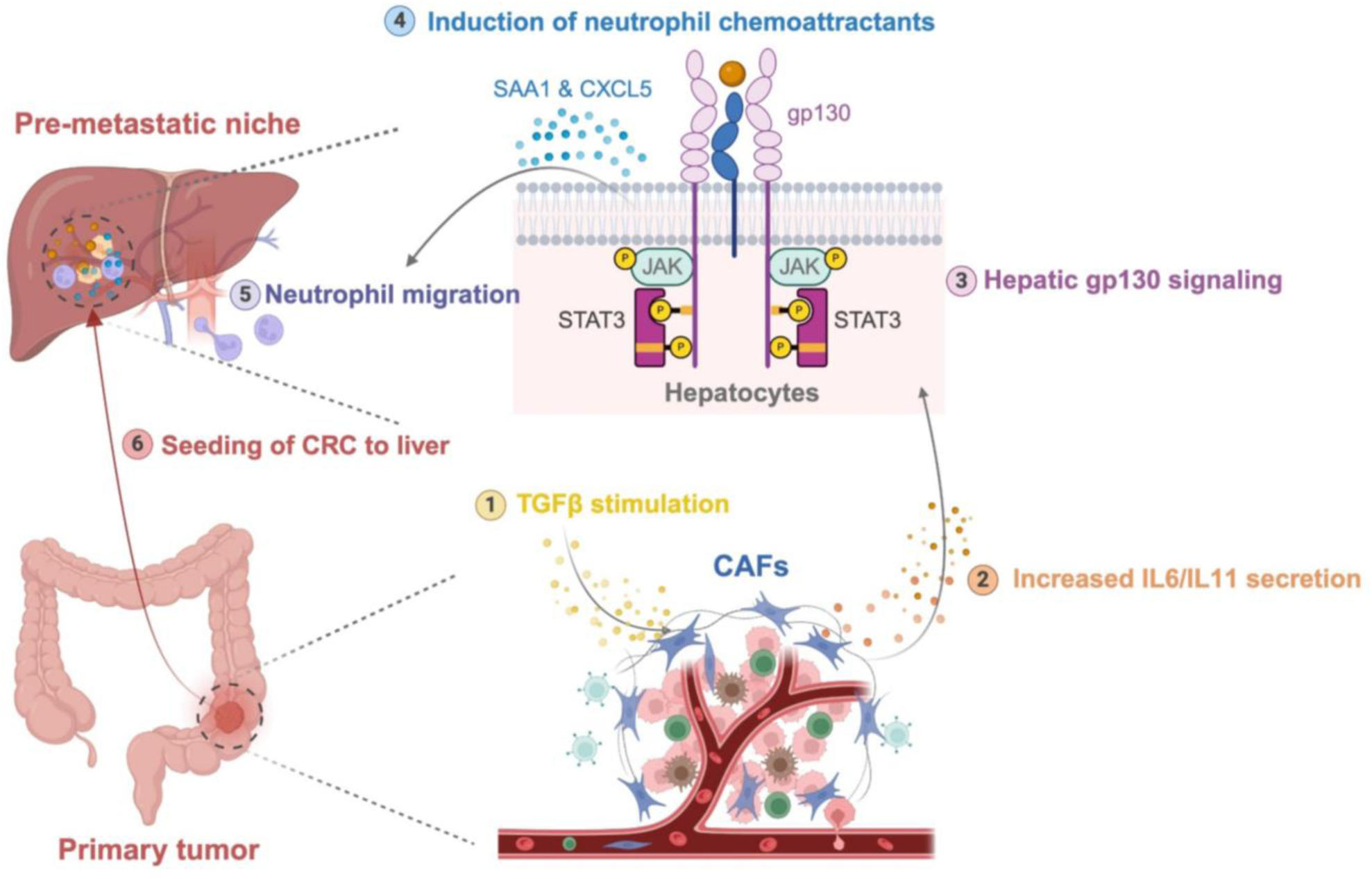
Schematic overview of molecular mechanisms driving hepatic metastasis in CMS4 CRC.TGFβ stimulation on primary CRC CAFs induces the secretion of IL-6 family members, notably IL-6 and IL-11. Subsequently, these CAF-derived IL-6 and IL-11 activate a gp130-dependent STAT3 signaling pathway in hepatocytes. This, in turn, leads to the production of neutrophil chemoattractants such as SAA1 and CXCL5, facilitating the migration of pro-metastatic neutrophils toward the liver. Ultimately, this signaling cascade shapes a pre-metastatic hepatic milieu favorable for the seeding of primary CRC tumors.

## Discussion

In our study, we identified TGFβ as a key driver for increased IL-6 expression specifically in CRC-derived CAFs. Under basal conditions, IL-6 expression in matched normal colon fibroblasts and CRC CAFs did not differ, emphasizing the influence of the TME in which these fibroblasts reside and the role of TGFβ expression/activation in the upregulation of IL-6. Given the high expression of TGFβ isoforms in the TME of CMS4 subtype CRC^3^, our findings reveal that all three isoforms can regulate IL-6, suggesting that CAFs can become active IL-6 producers in the context of the TGFβ-rich CMS4 TME. This increased IL-6 production by CAFs induces a pro-inflammatory program in hepatocytes when exposed to the IL-6 present in the supernatant produced by CAFs. This aligns with previous research highlighting the impact of primary tumor-derived cues on metastasis formation. While primary tumor-derived IL-6 has been shown to be essential for the formation of a hepatic pre-metastatic niche in a murine model of PDAC^19^, our study shows that IL-6 is only partially responsible for the increase in neutrophil chemoattractants SAA1 and CXCL5, with full dependence on the IL-6 co-receptor, gp130. Therefore, our focus extended to the broader IL-6 family of cytokines, in particular IL-11 and LIF that are also secreted in a TGFβ-regulated way by CAFs. Recent work in PDAC and rectal cancer has shown that a particular subset of CAFs, inflammatory CAFs (iCAFs), is characterized by the expression of these three IL-6 cytokine family members^41, 42^. Moreover, iCAFs have been associated with poor prognosis in rectal cancer and can confer resistance to neoadjuvant radiotherapy^42^. Our data suggest a potential role for TGFβ in modulating the expression of IL-6 family cytokines, which are correlated with unfavorable outcomes. Intriguingly, TGFβ has been described as a major molecular driver of the myofibroblastic CAF (MyCAF)^41, 43^ subset that have been shown to play a protective role by restraining CRC growth^44, 45^. Although MyCAFs are abundantly present in CMS4 CRC, this subtype demonstrates an aggressive rather than restrained phenotype, featuring the highest hepatic metastasis rate. These findings underscore the pleiotropic nature of TGFβ and its ability to regulate genes associated with both the MyCAF (αSMA) and iCAF (IL-6, IL-11, LIF) phenotype, further supporting the concept that CAF subset differentiation should be viewed as plastic and, potentially, reversible^41^.

We show that TGFβ-driven secretion of IL-6 family members in primary CRC CAFs further activates a gp130-dependent pro-inflammatory program in hepatocytes, leading to the upregulation of the neutrophil chemoattractants SAA1 and CXCL5. Similar to our findings, stromal cell-produced IL-6 in a mouse primary pancreatic tumor has been shown to activate STAT3 signaling in hepatocytes, significantly increasing the production of SAA1/2 and promoting myeloid cell accumulation and pro-metastatic niche formation^19^. Notably, the upregulation of SAAs by hepatocytes can also be detected in both pancreatic and colorectal cancer patients with liver metastases. In line with this, we observed a reduction in serum SAA1 levels after resection of the primary CRC in patients. Traditionally viewed as the first line of defense in the immune system, neutrophils are now increasingly recognized for their pro-metastatic role in the context of cancer^18, 23, 46, 47^. In this regard, it is worth noting that in the context of the KPN GEMM mouse model, used as a surrogate of the CMS4 subtype in mouse studies^33^, it has been previously reported that hepatic metastasis formation was neutrophil-dependent. Moreover, epithelial-derived TGFβ2 was required for the observed liver neutrophilia and this could implicate that increased expression of members of the IL-6 cytokine family, through TGFβ2 stimulation of CAFs, is also an intermediate step in this process. Using this model, we revealed that IL6-JAK-STAT signaling is indeed considerably upregulated in primary KPN tumors and that an increase in liver neutrophilia is already observed at the pre-metastatic stage. Furthermore, our data also suggest that CAFs in liver metastases still retain the ability to increase IL-6 family of cytokine expression upon exposure to TGFβ, suggesting that this mechanism may further support the metastatic niche once it has been formed.

In conclusion, our data reveal that TGFβ signaling in CAFs actively contributes to the formation of a pre-metastatic hepatic niche and that this mechanism might play a role in CMS4 subtype CRC metastasis formation and progression. Moreover, these findings support exploring the use of already clinically approved TGFβ^16, 48^ and IL-6 family-gp130-JAK axis^49–51^ inhibitors as means of chemoprevention of hepatic metastasis, specifically in patients with CMS4 CRC, by disrupting this pro-metastatic CAF-hepatocyte-neutrophil axis.

## Acknowledgements

We thank Eveline de Jonge-Muller, Léonie Plug, Michael Gillespie and Megan Mills for technical support. We thank the Flow cytometry Core Facility (FCF) of Leiden University Medical Center (LUMC) in Leiden, the Netherlands, for technical support and cell sorting assistance. We thank Dr Catherine Winchester of the CRUK Scotland Institute for review of the manuscript. Overview figures were created with BioRender.com.

## Author contributions

T.J.H. and S.A. performed experiments and associated data analysis. E.v.d.W, I.S. and S.G.T.J. performed experiments. M.W, T.R.M.L., and O.J.S. performed murine experiments and associated data analysis. K.J.L. performed analyses on the CMS dataset. E.M.E.V. and L.J.A.C.H. supervised and assisted in data interpretation. T.J.H., S.A., E.M.E.V, and L.J.A.C.H. co-wrote the paper. All authors contributed to the review and editing of the final manuscript.

## Funding

TJ Harryvan is sponsored by a personal PhD grant from the Leiden University Medical Center. EME Verdegaal is sponsored by a grant from the Dutch Cancer Society-KWF grant 2021-13673. I Stouten is supported by a grant from the Bontius Foundation. OJ Sansom and M White are funded by Cancer Research UK (A31287 core funding to the CRUK Scotland Institute and DRCQQR-May21\100002 core program funding to OJ Sansom).

## Data availability

The data that support the findings of this study are available from the corresponding author upon reasonable request.

## Competing interests

None declared.

## Notes

### Competing Interest Statement

The authors have declared no competing interest.

