## Supplementary Information for "TGFβ signaling in cancer-associated fibroblasts drives a hepatic gp130-dependent pro-metastatic inflammatory program in CMS4 colorectal cancer subtype"

**Figure legends**

***Supplementary figure 1.*** **BMP6 expression in CAFs and tissues. A.** *BMP6* RNA expression in primary CRC CAFs (N=1) or the CCD-18Co colon fibroblast line (N=1) after stimulation with TGFβ1. **B.** BMP6- protein expression in lysates of normal colon tissue (N=10, healthy donors) and primary CRC (N=10, patients), corrected for total protein content.

***Supplementary figure 2.*** **GSE39394 data on TGFβ stimulation of CCD-18Co colon fibroblasts.** RNA-sequencing of the normal colon fibroblast cell line CCD-18Co GSE39394 dataset. Red and blue circles indicate up and downregulated genes, respectively. The location of *IL6*, *IL11* and *LIF* is shown in green circles.

***Supplementary figure 3.*** **Neutrophil chemoattractant expression in hepatocytes upon CAF priming. A.** Expression of indicated neutrophil chemoattractants after stimulation of wildtype Huh-7 cells with DMEM 0% FCS (Control), IL-6 (50ng/ml), CAF CM (N=3) or TGFβ1-stimulated CAF CM (N=3). ** $p \leq 0.01$, * $p \leq 0.05$ determined by one-way ANOVA with correction for multiple testing (Dunnett's test). **B.** Schematic overview of the IL-6 signaling pathway and the levels of intervention with either blocking antibodies (siltuximab, tocilizumab, anti-gp130) or JAK inhibitor (tofacitinib).

***Supplementary figure 4.*** **Generation and verification of gp130 knockdown and knock-out Huh7 cells. A.** RNA expression of *gp130* in Huh-7 cells lentivirally transduced with a non-mammalian targeting ($SH^{Ctrl}$) or one of 4 different gp130 knockdown constructs. **B.** Western blot of $SH^{Ctrl}$ and $gp130^{KD}$ Huh-7 cell lines stained for gp130, pSTAT3. β-actin is used as loading control. Cells were either treated with DMEM 0% FCS (Control) or IL-6 (50ng/ml) to study gp130-dependent signaling. **C.** Western blot of $SH^{Ctrl}$ and $gp130^{KD}$ Huh-7 cell lines ($gp130^{KD1}$ & $gp130^{KD4}$) with and without stimulation with TGFβ1 CAF CM. **D.** FACS verification of $gp130^{KO}$ clonal Huh-7 cell line. **E.** PCR on cDNA from Huh-7 vector control, $gp130^{KO}$ and $gp130^{KO+Rescue}$ showing the absence or presence of gp130 or gp130-IRES amplicon. β-actin was used as a reference gene. **F.** Sanger sequence confirmation of generation of $gp130^{CA}$ plasmid (bottom sequence) from the wildtype pLV-THS-CMV-*gp130*-neomycin vector (upper sequence).

***Supplementary figure 5.*** **IL-11 contributes to the hepatic gp130-mediated inflammatory program. A.** RNA expression of members of IL-6 family of cytokines in CRC CAFs upon TGFβ1 stimulation (N=5). ND: non detectable. qPCR values have been log10 transformed. **** $p \leqslant 0.0001$, *** $p \leqslant 0.001$ determined by two-way ANOVA with correction for multiple testing. **B.** RNA expression of *IL11* and *LIF* in CRC CAFs stimulated with TGFβ1. N=2 or N=3 is used per CAF, independent biological experiments **** $p \leqslant 0.0001$determined by two-way ANOVA with correction for multiple testing. **C.** RNA expression of *IL11RA* in Huh-7 cells lentivirally transduced with a non-mammalian targeting ($SH^{Ctrl}$) or two different IL11RA knockdown constructs (N=1). **D.** Western blot for pSTAT3 of CAF-primed Huh-7 cells that were lentivirally transduced with $SH^{Ctrl}$ or one of 2 *IL11RA* knockdown constructs. β-actin is used as loading control.

***Supplementary figure 6.*** **RNA expression of members of IL-6 family of cytokines.** RNA expression of *CTF1* and *CLCF1* in CRC CAFs upon TGFβ1 stimulation. qPCR values have been log10 transformed. **** $p \leqslant 0.0001$, *** $p \leqslant 0.001$, * $p \leqslant 0.05$ determined by two-way ANOVA with correction for multiple testing.

***Supplementary figure 7.*** **Organoid vs. tumor tissue expression of IL-6 family of cytokines in KPN tumor. A-C**. Normalized counts of *IL6* (**A**), *IL11* (**B**) and *LIF* (**C**) expression in 1 KPN organoid line and whole tumor tissue generated through KPN-organoid injection into 3 mice.

### Supplementary figure 1

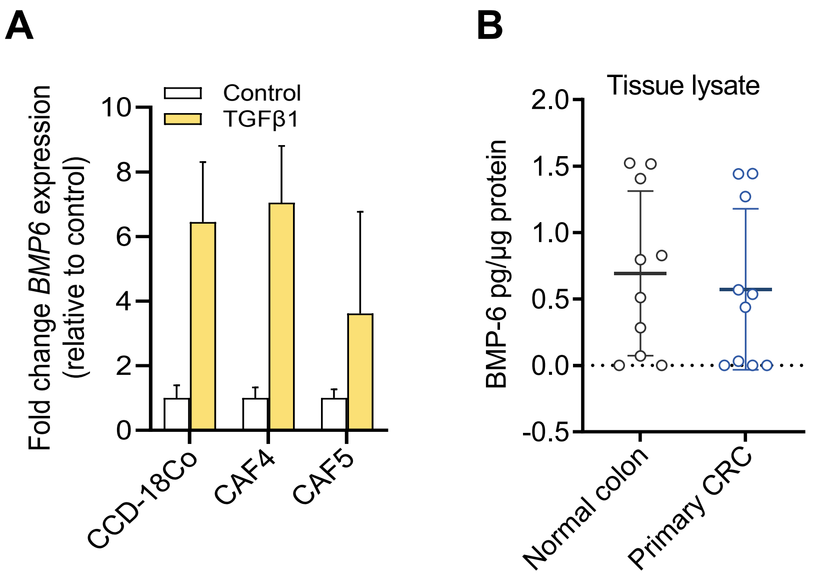

### Supplementary figure 2

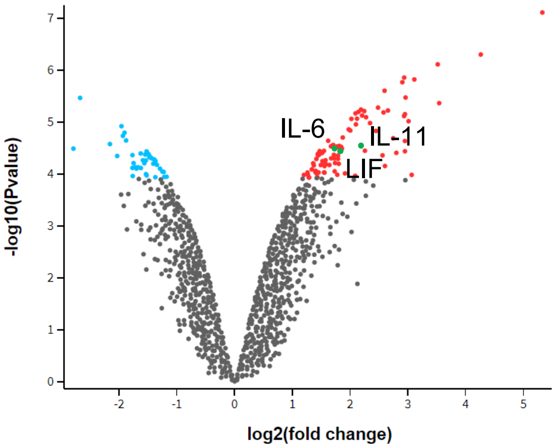

### Supplementary figure 3

**A**

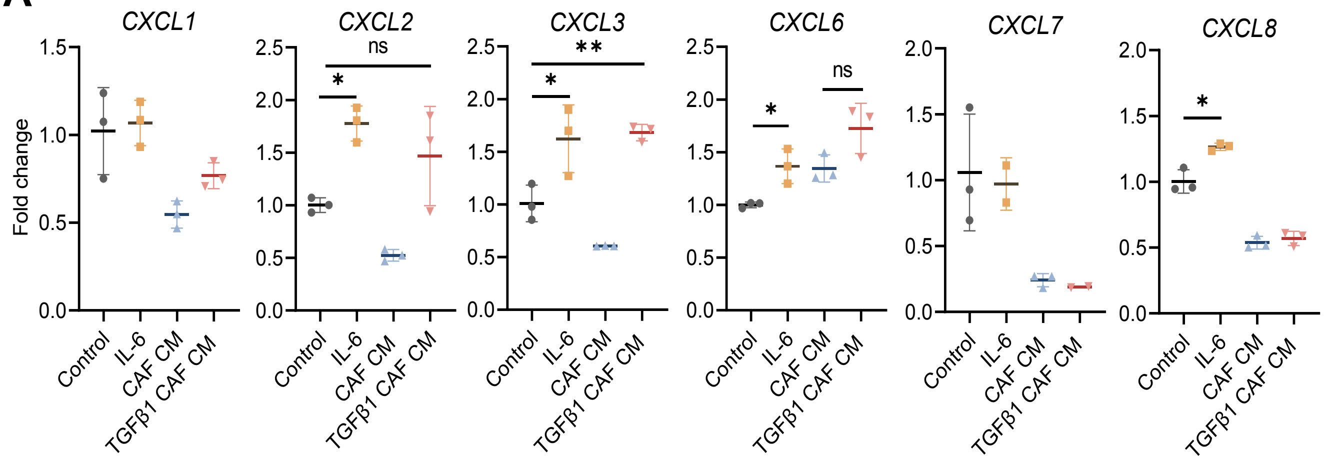


**B**

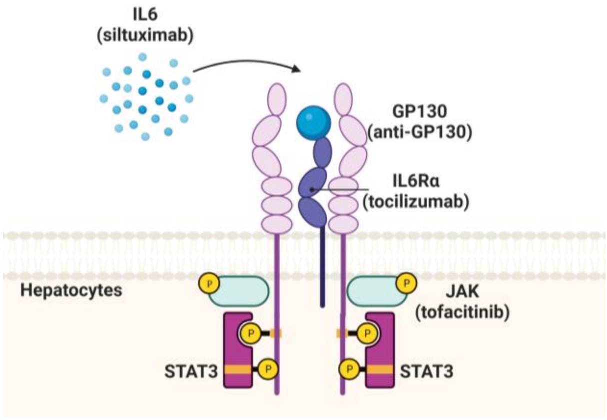

### Supplementary figure 4

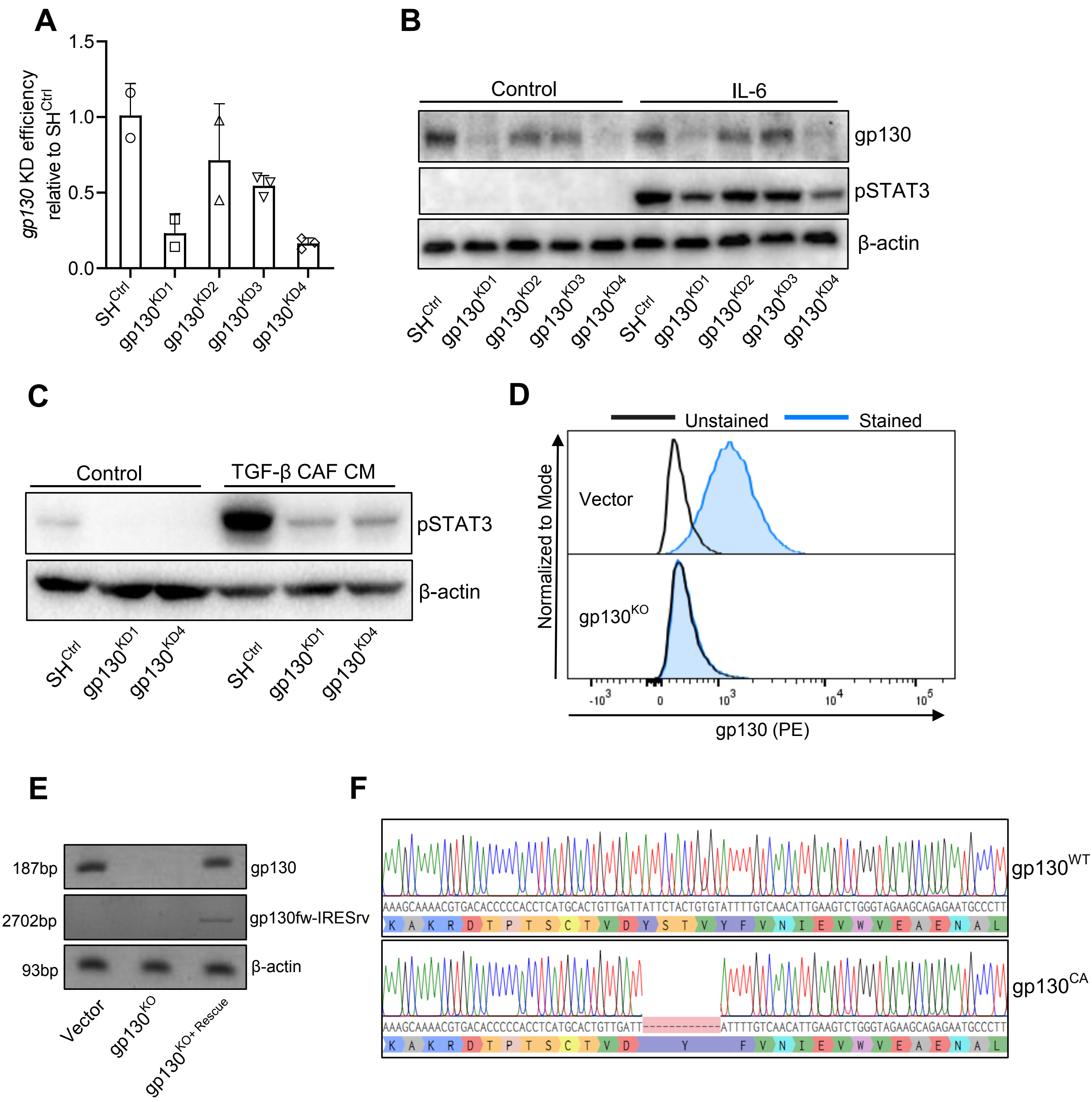

### Supplementary figure 5

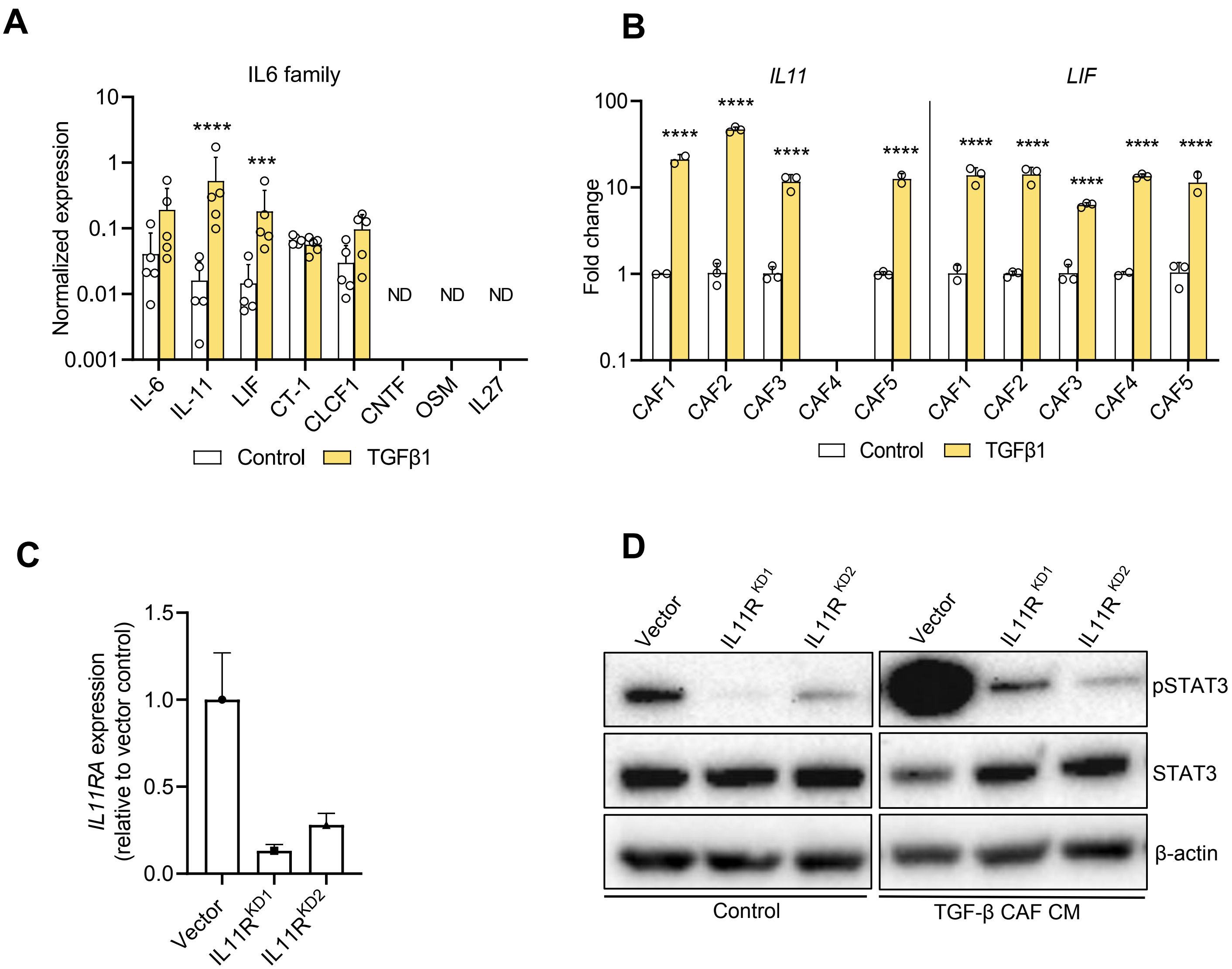

### Supplementary figure 6

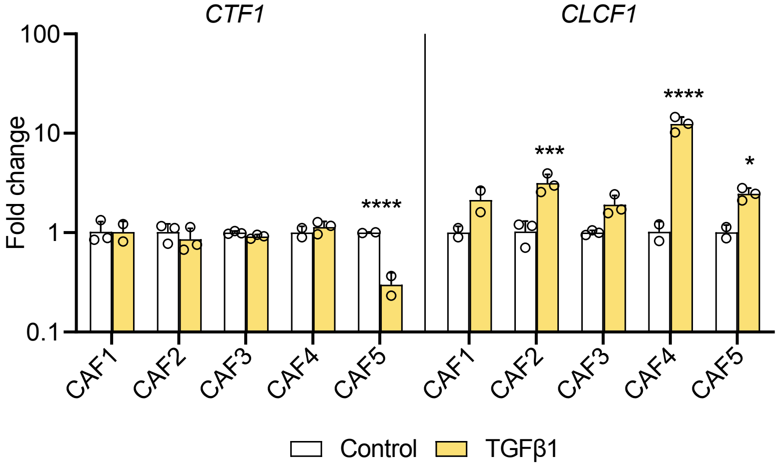

Supplementary figure 7

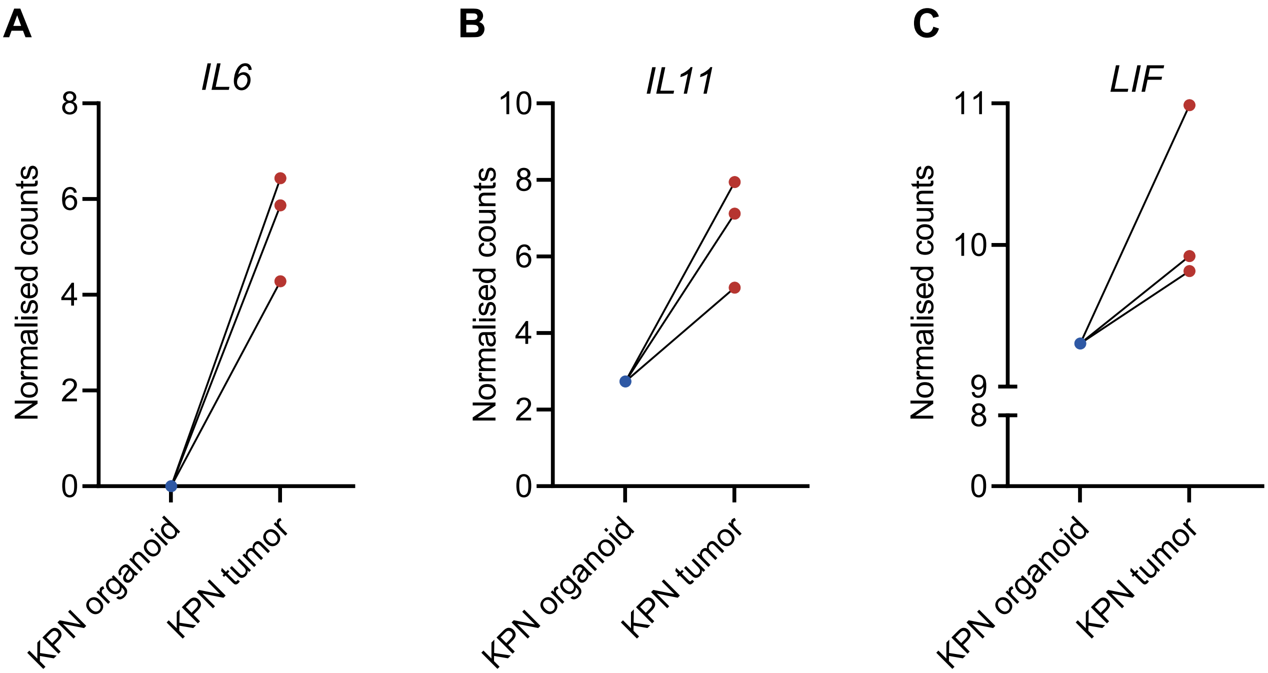

**Supplementary table 1. Mission shRNA constructs**

| Mission shRNA constructs | Sequence (5'-3') |
|---|---|
| Human gp130 KD#1 (TRCN0000058284) | CCAGTCCAGATATTTCACATT |
| Human gp130 KD#2 (TRCN0000058285) | CCCATACTCAAGGCTACAGAA |
| Human gp130 KD#3 (TRCN0000058287) | CGGCCAGAAGATCTACAATTA |
| Human gp130 KD#4 (TRCN0000289773) | CCAGTCCAGATATTTCACATT |
| Human IL11RA KD#1 (TRCN0000058934) | GCTCAAGTTCCGTTTGCAGTA |
| Human IL11RA KD#2 (TRCN0000058935) | CCCAGCCAGATCAGCGGTTTA |

**Supplementary table 2. Primer sequences**

| Gene | Primer sequence (5'-3') |
|---|---|
| *CXCL1* | Fw: CAGAAGGGAGGAGGAAGCTC<br>Rv: CTCTGCAGCTGTGTCTCTCT |
| *CXCL2* | Fw: CGCCCAAACCGAAGTCATAG<br>Rv: AGCTTCCTCCTTCCTTCTGG |
| *CXCL3* | Fw: TTCACCTCAAGAACATCCAAAGTG<br>Rv: TTCTTCCCATTCTTGAGTGTGGC |
| *CXCL5* | Fw: CAGACCACGCAAGGAGTTCATC<br>Rv: TTCCTTCCCGTTCTTCAGGGAG |
| *CXCL6* | Fw: CGCTGGTCCTGTCTCTGCT<br>Rv: GTTTTTCTTGTTTCCACTGTCC |
| *CXCL7* | Fw: TGCTCTGGCTTCCTCCACCAAA<br>Rv: ACACATGCAGCGGAGTTCAGCA |
| *SAA1* | Fw: AGGCTCAGACAAATACTTCCATGC<br>Rv: TCTCTGGCATCGCTGATCACTTCT |
| *IL8* | Fw: TCCTGATTTCTGCAGCTCTGT<br>Rv: AAATTTGGGGTGGAAAGGTT |
| *gp130* | Fw: ACATTCGGACAGCTTGAACAG<br>Rv: ACTTGTGTGTTGCCCATTCAGAT |
| *IL11RA* | Fw: GCTGTGTTGTCCTGGAGTGACT<br>Rv: ATGTAGGTGCCCTCATCAGTGC |

| | |
|---|---|
| *ACTB* | Fw: GTTGTCGACGACGAGCG<br>Rv: GCACAGAGCCTCGCCTT |
| *HPRT* | Fw: CCTAAGATGAGCGCAAGTTGAA<br>Rv: CCACAGGACTAGAACACCTGCTAA |
| *B2M* | Fw: TGCTGTCTCCATGTTTGATGTATCT<br>Rv: TCTCTGCTCCCCACCTCTAAGT |
| *IL6* | Fw: AGTGAGGAACAAGCCAGAGC<br>Rv: GTCAGGGGTGGTTATTGCAT |
| *CT1* | Fw: ATCCGTCAGACACACAGCCTTG<br>Rv: GAGAAGCTGGGCAGCCCGAAG |
| *CNTF* | Fw: TCAGACCTGACTGCTCTTACGG<br>Rv: TTGGAGTCGCTCTGCCTCGGT |
| *OSM* | Fw: GAAAGAGTACCGCGTGCTCCTT<br>Rv: CTCTCAGTTTAGGAACATCCAGG |
| *IL27* | Fw: GGAATCTCACCTGCCAGGAGTG<br>Rv: TGGTGGAGATGAAGCAGAGACG |
| *LIF* | Fw: AGATCAGGAGCCAACTGGCACA<br>Rv: GCCACATAGCTTGTCCAGGTTG |
| *IL11* | Fw: GGACCACAACCTGGATTCCCTG<br>Rv: AGTAGGTCCGCTCGCAGCCTT |
| *CLCF1* | Fw: CCGCTACCTGGAGCACCAACT<br>Rv: CACACCTCCAAGTCAACAGTGG |
| *IL6R* | Fw: ACGCCTTGGACAGAATCCAG<br>Rv: GAATCTTGCACTGGGAGGCT |
| *gp130* | Fw: GAATCTTGCACTGGGAGGCT<br>Rv: ACATTCGGACAGCTTGAACAG |
| *IL11R* | Fw: GCTGTGTTGTCCTGGAGTGACT<br>Rv: ATGTAGGTGCCCTCATCAGTGC |
| *IRES rv* | CCTCACATTGCCAAAAGACG |

1
